# Hacking brain development to test models of sensory coding

**DOI:** 10.1101/2023.01.25.525425

**Authors:** Maria Ahmed, Adithya E. Rajagopalan, Yijie Pan, Ye Li, Donnell L. Williams, Erik A. Pedersen, Manav Thakral, Angelica Previero, Kari C. Close, Christina P. Christoforou, Dawen Cai, Glenn C. Turner, E. Josephine Clowney

## Abstract

Animals can discriminate myriad sensory stimuli but can also generalize from learned experience. You can probably distinguish the favorite teas of your colleagues while still recognizing that all tea pales in comparison to coffee. Tradeoffs between detection, discrimination, and generalization are inherent at every layer of sensory processing. During development, specific quantitative parameters are wired into perceptual circuits and set the playing field on which plasticity mechanisms play out. A primary goal of systems neuroscience is to understand how material properties of a circuit define the logical operations— computations--that it makes, and what good these computations are for survival. A cardinal method in biology—and the mechanism of evolution--is to change a unit or variable within a system and ask how this affects organismal function. Here, we make use of our knowledge of developmental wiring mechanisms to modify hard-wired circuit parameters in the *Drosophila melanogaster* mushroom body and assess the functional and behavioral consequences. By altering the number of expansion layer neurons (Kenyon cells) and their dendritic complexity, we find that input number, but not cell number, tunes odor selectivity. Simple odor discrimination performance is maintained when Kenyon cell number is reduced and augmented by Kenyon cell expansion.

## Introduction

In diverse bilaterians, chemosensory information is processed through parallel circuits that support innate versus learned interpretations (Ghosh et al., 2011; Marin et al., 2002; Miyamichi et al., 2011; Sosulski et al., 2011; Tanaka et al., 2004; Wong et al., 2002). Circuits for innate processing rely on developmental specification of distinct cell types that wire together in stereotyped patterns to connect sensory inputs to evolutionarily-selected behavioral responses (Chin et al., 2018; Clowney et al., 2015; Fişek and Wilson, 2014; Jefferis et al., 2007; Kobayakawa et al., 2007; Lin et al., 2011; Root et al., 2014; Troemel et al., 1997; Wang et al., 2018). In contrast, regions devoted to learned interpretation appear more like *in silico* computers, with the same circuit motif repeated thousands or millions of times (Albus, 1971; Ito, 1972; Marr, 1969; Minsky, 1952, n.d.). Such repetitive organization allows circuits for learned interpretation to function like switchboards, with the potential to connect any possible sensory representation (the caller) to any possible behavioral output (the receiver). Development of learning regions involves the specification of large groups of neurons with the same identity that are capable of receiving broad sensory inputs and that connect with neurons driving multiple potential behavioral outputs (Luo, 2021).

The quantitative wiring parameters that set up an organism’s potential to recognize stimuli and learn their meanings are dictated by the developmental identities of the neurons that comprise learning circuits. The transformation a neuron makes from input to output—its computation— depends on the structure of its wiring in the circuit and on its electrophysiological properties.

What can an animal even sense? What sorts of stimuli can it discriminate from one another? Can it extract general features from different contexts? How an animal perceives arbitrary stimuli--those whose meanings are not inscribed in the genome—and what it can learn about them depends on the architectural and physiological specifics of its associative learning circuits.

An “expansion layer” is a common motif observed in associative learning circuits where neurons receiving information about a set of sensory channels connect combinatorially onto a much larger set of postsynaptic cells (Albus, 1971; Ito, 1972; Marr, 1969). These layers are found in each of the major clades of animals with centralized brains and include the chordate pallium, cerebellum, and hippocampus; the arthropod mushroom body; and the cephalopod parallel lobe system. Beginning with Marr-Albus theory of the cerebellum in the 1970’s, expansion coding has been hypothesized to perform pattern separation. The ratio between sensory channels and expansion layer neurons, and the number of sensory inputs that individual expansion layer neurons receive, are theorized to be key parameters in balancing perception, discrimination, and generalization (Albus, 1971; Babadi and Sompolinsky, 2014; Cayco-Gajic and Silver, 2019; Hiratani and Latham, 2022; Ito, 1972; Jefferis et al., 2007; Jortner et al., 2007; Litwin-Kumar et al., 2017; Luo et al., 2010; Marr, 1969; Modi et al., 2020; Rajagopalan and Assisi, 2020). However, the perceptual and behavioral effects of altering hard-wired quantitative relationships have not been experimentally tested.

To test the functional utility of observed neural circuit structures, we initiate here a project of developmental circuit hacking. Using our knowledge of mushroom body structure and development in *Drosophila melanogaster*, we change quantitative relationships between presynaptic olfactory projection neurons and postsynaptic Kenyon cells *in vivo* (Elkahlah et al., 2020; Puñal et al., 2021). We then test perceptual and behavioral capabilities of these hacked-circuit animals.

The arthropod mushroom body is a facile model for testing how the architectural features of expansion layer circuits enable sensory representation and learned associations. Kenyon cells are the expansion layer neurons of the mushroom body; they receive input from diverse olfactory projection neurons (PNs) in the mushroom body calyx and send outputs to varied and broadly ramifying mushroom body output neurons (MBONs) in the mushroom body lobes (Aso et al., 2014a, 2014b). Uniglomerular PNs each receive input from a single kind of olfactory sensory neuron in one of 52 antennal lobe glomeruli (Grabe et al., 2016). These 52 types of PNs form multisynaptic boutons in the calyx (Butcher et al., 2012; Leiss et al., 2009; Yang et al., 2022). Each Kenyon cell has an average of 5-6 discrete dendritic “claws” that each innervate a single bouton; the cell therefore receives combinatorial olfactory input (Caron et al., 2013; Zheng et al., 2018). The sets of odor inputs to individual Kenyon cells are diverse and approximate a random sampling of available PN boutons (Caron et al., 2013; Eichler et al., 2017; Zheng et al., 2022). Kenyon cells typically require multiple active inputs to fire and therefore act as coincidence detectors with the potential to expand the animal’s perception from single channels to mixtures (Groschner et al., 2018; Gruntman and Turner, 2013; Li et al., 2013; Lin et al., 2014). Dopamine-mediated plasticity at Kenyon cell:MBON synapses in the lobes allows animals to learn associations between odors and temporally linked events (Cohn et al., 2015; Hige et al., 2015; Owald et al., 2015). Learning-dependent changes to animal behavior can endure for days (Tully et al., 1994).

In previous work, we developed methods to increase and decrease the number of Kenyon cells. We found that individual Kenyon cells received the same number of sensory inputs in these conditions, because presynaptic olfactory PNs adjusted their output repertoire as the number of Kenyon cells changed (Elkahlah et al., 2020). This developmental rule means that the size of the Kenyon cell population and the number of sensory inputs to individual Kenyon cell can be “programmed” as independent variables during development. Here, we develop methods to increase and decrease Kenyon cell claw number and therefore the number of olfactory inputs that individual Kenyon cells receive. We then use these circuit-hacked animals to test the effects of altering Kenyon cell number and claw number on sensory representations and associative learning behavior. We find that changing Kenyon cell number only modestly affects population-level odor responses. In contrast, Kenyon cell odor responses change bidirectionally as we change Kenyon cell dendritic claw number, such that Kenyon cells become less odor-selective as their input number grows. Remarkably, animals with reduced Kenyon cell population size can learn simple olfactory associations, and animals with augmented sets of Kenyon cells show improved associative learning. These results illuminate surprising functional and developmental principles and provide a novel method for testing the “purpose” of observed learning circuit architectures.

## Results

### Sparse odor coding is preserved despite perturbations to Kenyon cell numbers

There are ∼2000 Kenyon cells within the *Drosophila melanogaster* mushroom body that receive combinatorial inputs from ∼160 olfactory PNs (Figure 1A, B) (Aso et al., 2009; Bates et al., 2020; Li et al., 2020; Schlegel et al., 2021; Takemura et al., 2017). To determine the relationship between circuit form and function *in vivo*, we previously developed methods to change the number of Kenyon cells, altering the PN:Kenyon cell ratio (Elkahlah et al., 2020). When we varied the number of Kenyon cells from 500 to 4000 per hemisphere, we found that individual Kenyon cells retained their typical claw numbers—about six per cell--while PNs scaled their bouton numbers to suit Kenyon cell input demands. In mushroom bodies with a reduced repertoire of Kenyon cells, we observed that Kenyon cell odor responses remained selective and sparse—different populations of cells responded to different odors, and most cells responded to a minority of odors.

**Figure 1.**
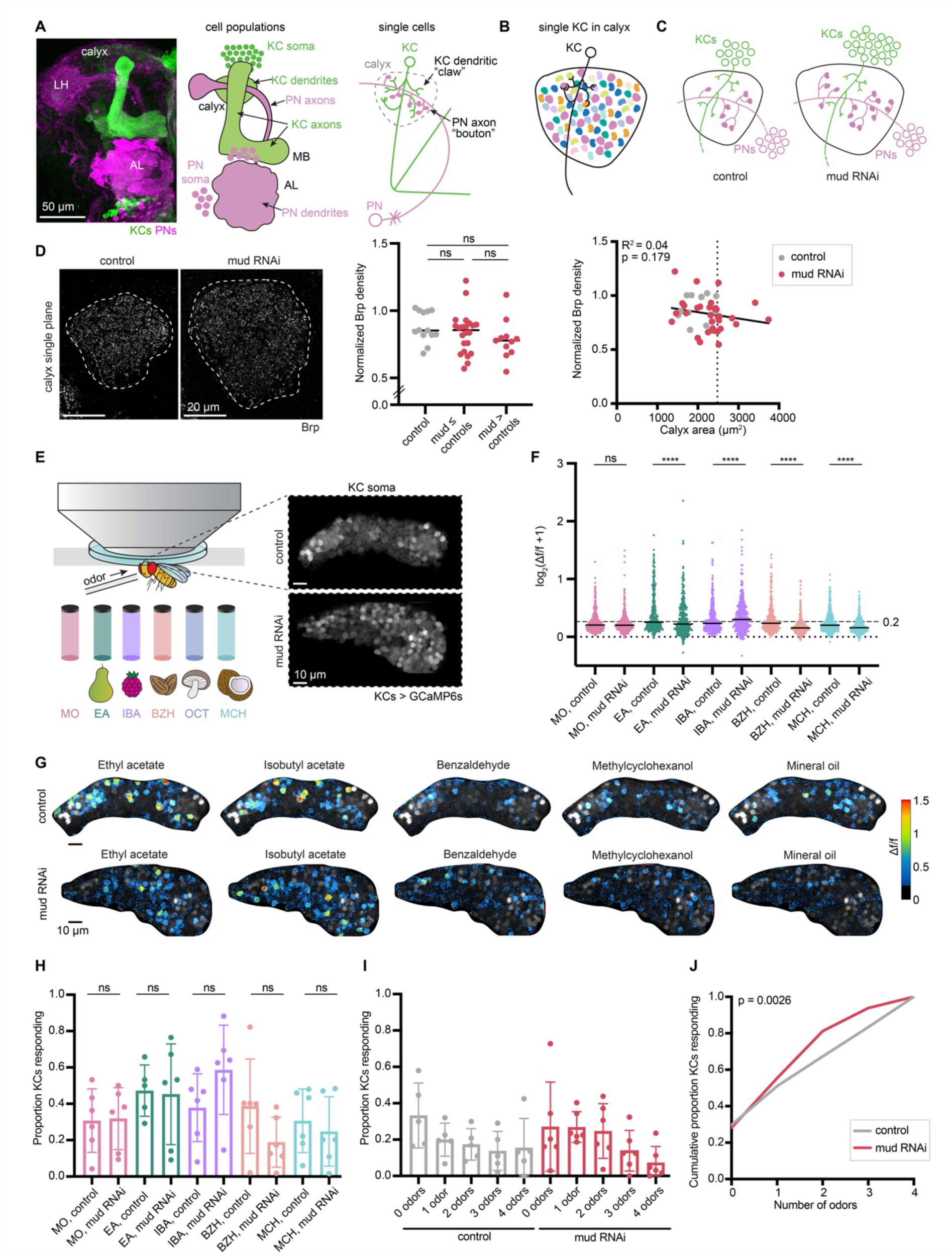
Sparse odor coding is preserved when Kenyon cell numbers are increased. (A) Image (left) and models (right) of olfactory projection neurons (PNs) and Kenyon cells (KCs) in the adult fly brain at the population level and single cell level. PNs (pink) receive input from olfactory sensory neurons in the antennal lobe (AL) and project to the mushroom body calyx and the lateral horn (LH). In the calyx, KCs (green) form dendritic “claws” that grab onto output sites of PNs called axon “boutons”. (B) An individual KC receives input from an average of 5-6 PNs. The different colors of boutons indicate PN types receiving input from different glomeruli. (C) Model of the effect of increasing KC numbers on calyx development. PNs increase bouton numbers to match the increase in KCs, while KCs maintain the same number of dendritic claws per cell. PN number is unchanged. (D) Left: Confocal slice of Brp signal in control calyx and OK107>*mud* RNAi calyx; maximum cross-sectional area is circled. Middle: Quantification of Brp density, normalized to fluorescence in an unmanipulated brain region, the protocerebral bridge. Significance: unpaired t-test for each pairwise comparison. Right: The relation of normalized Brp density to maximum cross-sectional area of calyx. Dotted line represents the calyx area cut-off to define Kenyon cell-increased *mud* RNAi brains (“mud > controls”). Each data point represents a single hemisphere. (E) Schematic of preparation used for *in vivo* functional imaging in an adult fly. Odor vials for the mechanosensory control (mineral oil; MO) and odors used, along with their smells, are shown below: ethyl acetate (EA), isobutyl acetate (IBA), benzaldehyde (BZH), octanol (OCT), and methylcyclohexanol (MCH). Responses were imaged from KC soma by GCaMP6s expression; example images of a control KC soma cloud, and *mud* RNAi, expanded KC cloud are shown. (F) Peak odor responses of all cells plotted on a log scale, aggregated from all six analyzed hemispheres of each condition. Dashed line indicates 20% Δf/f threshold. Dotted line indicates Δf/f = 0. Black horizontal bars show medians. Significance: Mann-Whitney test between control and *mud* RNAi for each stimulus. Here and throughout, *: p<0.05, **: p<0.01, ***: p<0.001, ****: p<0.0001, ns: non-significant. n = 349-405 cells (control), n = 341 cells (*mud* RNAi) for each odor. (G) Representative images of KC somatic odor responses in control hemisphere and increased-KC *mud* RNAi hemisphere. Grayscale backdrop indicates the cells, Δf/f scale is shown such that all cells with responses < 0.2 are colored black, responses > 1.5 are colored red. See also Figure S1A. (H) Proportion of cells in each sample responding to each odor, and mineral oil above 0.2 Δf/f threshold. Significance: unpaired t-test. (I) For samples in which the same cells could be tracked across all odor presentations, proportion of cells responding to 0, 1, or multiple odors is shown. Bar plots in (H, I) show mean ± SD, and black circled points highlight the particular hemispheres shown in (G). See also Figure S1B. (J) Cumulative proportion of cells responding from 0 up to 4 odors. Lines represent mean of all control (gray; n = 349) and increased-KC *mud* RNAi cells (red; n = 341). Significance: Kruskal-Wallis test; K-S distance=0.139.

Next, we sought to ask how supernumerary Kenyon cells respond to odors. We have previously amplified Kenyon cell numbers by knocking down *mushroom body defect* (*mud*) in Kenyon cell neuroblasts (Elkahlah et al., 2020). Mud/NuMA is a spindle orientation protein that allows neuroblasts to divide asymmetrically; loss of *mud* results in occasional neuroblast duplication (Guan et al., 2000; Prokop and Technau, 1994; Siller et al., 2006). By driving UAS *mud* RNAi in Kenyon cell neuroblasts under control of OK107-Gal4, we can produce mushroom bodies with as many as 4000 cells (Elkahlah et al., 2020). As we described previously, OK107 labels both progenitors and differentiated Kenyon cells; however, *mud* is only transcribed in progenitors (Elkahlah et al., 2020).

When Kenyon cell numbers are increased, individual neurons retain claw numbers similar to wild type, while olfactory PNs show remarkable developmental plasticity by scaling up their bouton numbers to match the change in postsynaptic cell numbers (Figure 1C) (Elkahlah et al., 2020). This suggests that the number of synapses per Kenyon cell will remain unchanged. To ask if that is the case, we immunostained for an active zone marker, Bruchpilot (Brp), which accurately reflects electron microscopy synapse counts (Figure 1D) (Lillvis et al., 2022; Wagh et al., 2006). As neuroblast duplications are stochastic, the *mud* RNAi animals show considerable variability in Kenyon cell numbers. Therefore, we grouped *mud* RNAi animals into “mud ≤ controls” and “mud > controls” based on their maximum cross-sectional calyx area, which we have previously found to correlate with Kenyon cell numbers (Elkahlah et al., 2020).

Normalized Brp density did not vary with calyx area, suggesting that the number of input synapses per Kenyon cell is maintained in the increased-Kenyon cell calyces (Figure 1D).

To observe odor responses in brains with supernumerary Kenyon cells, we expressed GCaMP6s in all Kenyon cells and imaged somatic responses to odors in adult flies. Each fly was given a sequence of 4 different odors diluted 1:100 in mineral oil solvent and a mineral oil-only control, to capture motion and solvent responses (Figure 1E). In total, we imaged responses in 14 control hemispheres and 20 *mud* RNAi hemispheres. We observed robust odor responses in each condition, with only 1/14 control and 1/20 *mud* RNAi hemispheres completely unresponsive.

We selected *mud* RNAi data sets showing expanded calyces for analysis, as described previously. Each image was motion corrected in Suite2p; after motion correction, 6 control and 6 increased-Kenyon cell *mud* RNAi hemispheres were stable enough to follow individual cells over time (Pachitariu et al., 2017). Full inclusion criteria are described in the methods.

For each dataset here and throughout, we defined individual ROIs for cells that remained spatially stable across the session. We then calculated the peak fluorescence change (Δf/f) for each cell following delivery of each stimulus. We plotted the response magnitude for all cells pooled, to compare the aggregate distribution of responses to each stimulus (Figure 1F). Median odor responses were similar between control and increased-Kenyon cell calyces, though the shape of the distribution differed statistically (Figures 1F-G, S1A). We then chose a 20% increase in fluorescence from baseline level (Δf/f > 0.2) as the threshold for defining “responsive” cells based on bifurcation of the response distribution along this cutoff (Figure 1F).

We next quantified the proportion of Kenyon cells per calyx responding to each odor. The proportions were strikingly similar in the two conditions (Figure 1H), as we previously observed in reduced-Kenyon cell calyces (Elkahlah et al., 2020). In each condition, ethyl acetate and isobutyl acetate stimulated ∼40-60% of Kenyon cells, while benzaldehyde stimulated ∼20-40% of cells and methylcyclohexanol stimulated ∼30% of cells (Figure 1H). A discussion of baseline Kenyon cell odor response rates measured under different experimental conditions is provided in the methods and in (Elkahlah et al., 2020). The variability in responses to each odor across different animals is expected due to the stochastic nature of innervation of Kenyon cells by PNs (Caron et al., 2013).

Next, we asked how many odors each cell responded to and quantified the proportion of Kenyon cells per calyx responding to 0, 1, or multiple odors. In each condition, ∼50% of cells responded to 0 or 1 odor, and ∼10-15% of cells responded to all 4 odors (Figures 1I-J, S1B). Thus, populations of supernumerary Kenyon cells still respond sparsely to odors, and individual cells remain odor selective. Among expanded calyces, calyx size did not predict response sparseness (Figure S1C). Taken together, these results demonstrate that Kenyon cells maintain sparse odor coding even when the Kenyon cell population is increased.

### Knocking down *Tao* in Kenyon cells increases dendritic claws per cell

We hypothesize that the preservation of sparse odor responses in calyces with altered Kenyon cell number is due to maintenance of Kenyon cell dendritic claw number, and thus density of PN innervation, in those conditions. As we sought to test the influence of input density on the sparseness and selectivity of odor responses, we next searched for ways to directly alter the number of claws on each Kenyon cell. Tao is a kinase in the Mst/Ste20-family that interacts with par-1 to regulate microtubule dynamics and has recently been shown to negatively regulate dendritic branching (Dan et al., 2001; Hu et al., 2022, 2020; King et al., 2011; Mitsopoulos et al., 2003). We found that knockdown of *Tao* in all Kenyon cells under control of OK107 sharply expanded calyx size while preserving animal viability.

We hypothesized that calyx expansion was due to expansion of the dendritic arbors of individual cells. To test this, we developed an ultra-sparse version of the Bitbow labeling approach (Li et al., 2021; Veling et al., 2019). This Bitbow variant, mBitbow 2.2, relies on a double-recombinase approach and allows us to label as few as 1/1000 Gal4-positive cells (Figures 2A, 2B, S2). Consistent with the increased calyx size, we found that individual cells in the *Tao* knockdown had a 50% increase in claw number (Figure 2C-E). The dendrites also spread out further in the calyx as if “searching” for more connections, even occasionally going outside the calyx. These Kenyon cells also show greater diversity in dendritic morphology; for example, Kenyon cells occasionally had highly branched calycal processes that failed to form curved claw structures (Figures 2C, S2B). Additionally, we observed that Kenyon cells number actually decreased by 25% upon *Tao* knockdown, so the expansion of dendritic arbors is likely the dominant factor underlying calycal expansion (Figure 2E-G). Similar to previous work in *Tao* mutants and *Tao* RNAi in Kenyon cells, we also observed Kenyon cell axonal defects: some Kenyon cell axons appeared to be either missing or misdirected to the calyx (King et al., 2011). Incursion of axons into the calyx could further inflate our measurements of calyx area beyond the expansion in dendritic claw numbers.

**Figure 2.**
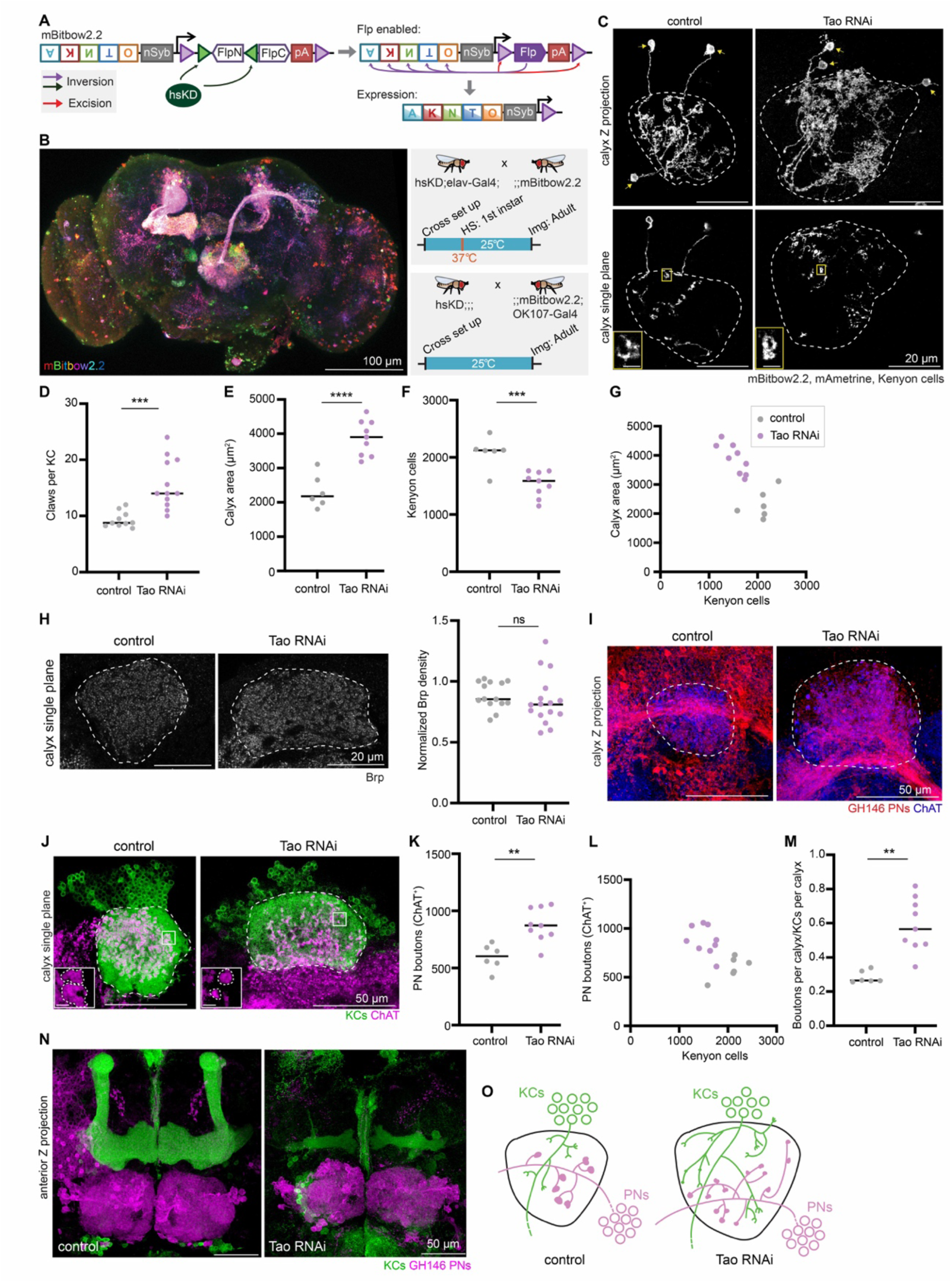
Knocking down *Tao* in Kenyon cells increases dendritic claws per cell and projection neuron bouton production. (A) Schematic of mBitbow2.2 design. mBitbow2.2 transgene contains two parts: a membrane-labeling mBitbow1.0 (Li et al., 2021) and a KD-controlled, self-excising flippase (KDonFlp) under the control of n-Synaptobrevin (nSyb) promoter. In the presence of KD, the N-terminal portion of flippase will be inverted to the correct orientation, hence enabling the expression of flippase under nSyb control in mature neurons. Similar to the mBitbow2.1 design, the flippase will initiate a Bitbow combination. Flippase can also excise itself, preventing sustained recombinations. The five fluorescent proteins shown are mAmetrine (A), tdKatushka2 (K), mNeonGreen (N), mTFP1 (T), and mKO2 (O). Detailed schematic is shown in Figure S2A. (B) Left: A representative adult brain Z-projection demonstrating dense mBitbow2.2 labeling in neurons. Top right: To produce dense labeling, progeny of the hsKD;elav-Gal4;; and ;;mBitbow2.2; flies are collected and heat shocked at 1^st^ instar larval stage at 37°C for 30 mins. Neurons labeled in diverse colors can be observed throughout the adult brain, with prominent neuropils evident, such as the mushroom body and projection neuron tracts from the antennal lobe. Bottom right: For sparse Kenyon cell labeling, hsKD;;; flies were crossed with ;;mBitbow2.2; OK107-Gal4. No heat shock was given. Crosses were housed at 25°C throughout. (C) Example images of sparsely labeled KCs in control and KC>*Tao* RNAi calyces: maximum Z-projection (top), single confocal slice (bottom). Dashed lines outline the calyx, and arrows mark the KC soma. In the examples shown, 3 KCs are labeled by mBitbow2.2 (mAmetrine shown). Insets zoom in on a single claw structure from the yellow boxed region in the corresponding image. Scale bar for inset: 2mm. See also Figure S2B. (D) Number of dendritic claws per Kenyon cell in control (gray) and KC>*Tao* RNAi (purple) hemispheres. Each data point represents a single hemisphere throughout this figure. (E-G) Maximum calyx cross-sectional area (E), number of KCs (F), and the relationship between KC number and maximum calyx cross-sectional area (G) in control (gray) and KC>*Tao* RNAi hemispheres (purple). (H) Left: Confocal slice showing Brp signal and maximum cross-sectional area (circled) of control and KC>*Tao* RNAi calyces. Right: Quantification of Brp density, normalized to fluorescence in an unaffected brain region, the protocerebral bridge. (I) Maximum intensity projection of confocal stack of calyx bouton production by GH146^+^ PNs (red) in control and KC>*Tao* RNAi hemispheres. ChAT immunostaining (blue) indicates calyx extent. (J) Confocal slice of control calyx and KC>*Tao* RNAi calyx. OK107 labels KCs (green) and ChAT labels projection neuron boutons (magenta). Representative images selected with median number of boutons from each condition, and image slice chosen to show the middle plane of boutons. Dashed lines outline the calyx. Inset zooms in on the bouton morphology differences from the boxed region in the image. Scale bar for inset: 2 µm. (K-M) PN bouton number (K, L), and ratio of ChAT^+^ boutons per calyx versus KCs per calyx (M) in control (gray) and KC>*Tao* RNAi hemispheres (purple). (N) Maximum intensity projection of confocal stacks taken from anterior side of control and KC>*Tao* RNAi brains. KCs (green) show mushroom body lobes. GH146^+^ PNs (magenta) show antennal lobe and PN cell bodies. (O) Model of the effect of increasing the number of dendritic claws per Kenyon cell. *Tao* RNAi KC dendrites spread out further than controls. PN bouton production increases, boutons appear smaller, and the PN tract shifts ventrally. Throughout the figure, significance: unpaired t-test, black horizontal bars represent median.

We next examined how *Tao* knockdown influenced innervation by the presynaptic boutons of PNs. We first examined overall presynaptic density by staining for Brp (Figure 2H). Despite the expansion of calyx size in these animals, Brp density did not change, suggesting that each Kenyon cell claw is innervated by the same number of presynaptic sites in control and *Tao* knockdown animals. We then examined how these synapses are organized into multisynaptic presynaptic terminals by counting the number of acetylcholine-positive boutons in each calyx (bouton counting procedure is described in the methods and demonstrated in (Elkahlah et al., 2020)). We observed an increased number of PN boutons per calyx (Figure 2I, K-M). Bouton morphology was also affected: boutons appeared smaller in their cross-sectional area in the *Tao* RNAi animals, suggesting that Kenyon cell claws might be shaping PN bouton structures (Figure 2J). Additionally, labeling 100 PNs with the GH146 driver highlighted a ventral shift of the PN axon tract in the calyx (Figure 2I). Gross antennal lobe morphology was unaffected (Figure 2N). Further description of these anatomic effects will be provided in a future publication.

In conclusion, knocking down *Tao* in Kenyon cells expanded calyx size and Kenyon cell dendritic claw number. Projection neurons produced an increased number of boutons that matched the increase in dendritic claws per Kenyon cell, which would likely result in an increase in odor inputs to each cell (Figure 2O).

### Odor selectivity is reduced in Kenyon cells with increased dendritic claw numbers

To test if increasing inputs to Kenyon cells alters combinatorial odor coding, we next expressed GCaMP6s under control of OK107 to measure odor responses of *Tao* knockdown Kenyon cells. Remarkably, we observed a strong increase in the number of cells responding to each odor and to mineral oil (Figures 3A, C, S3A). We found that twice as many Kenyon cells responded to each stimulus in the increased-dendrite condition as compared to the controls (e.g. ∼60% cells for methylcyclohexanol compared to ∼30% in controls) (Figure 3D). In some cases, we even observed all Kenyon cells responding to all stimuli (Figure 3A; red outlined points in Figure 3D, E). We hypothesized that with more cells responding to each stimulus, there would be more overlap in the representations of different stimuli. To test this, we measured Pearson correlation value between each stimulus; indeed, pairwise correlations in responses to each odor or mineral oil were higher in *Tao* knockdown animals (Figure 3B).

**Figure 3.**
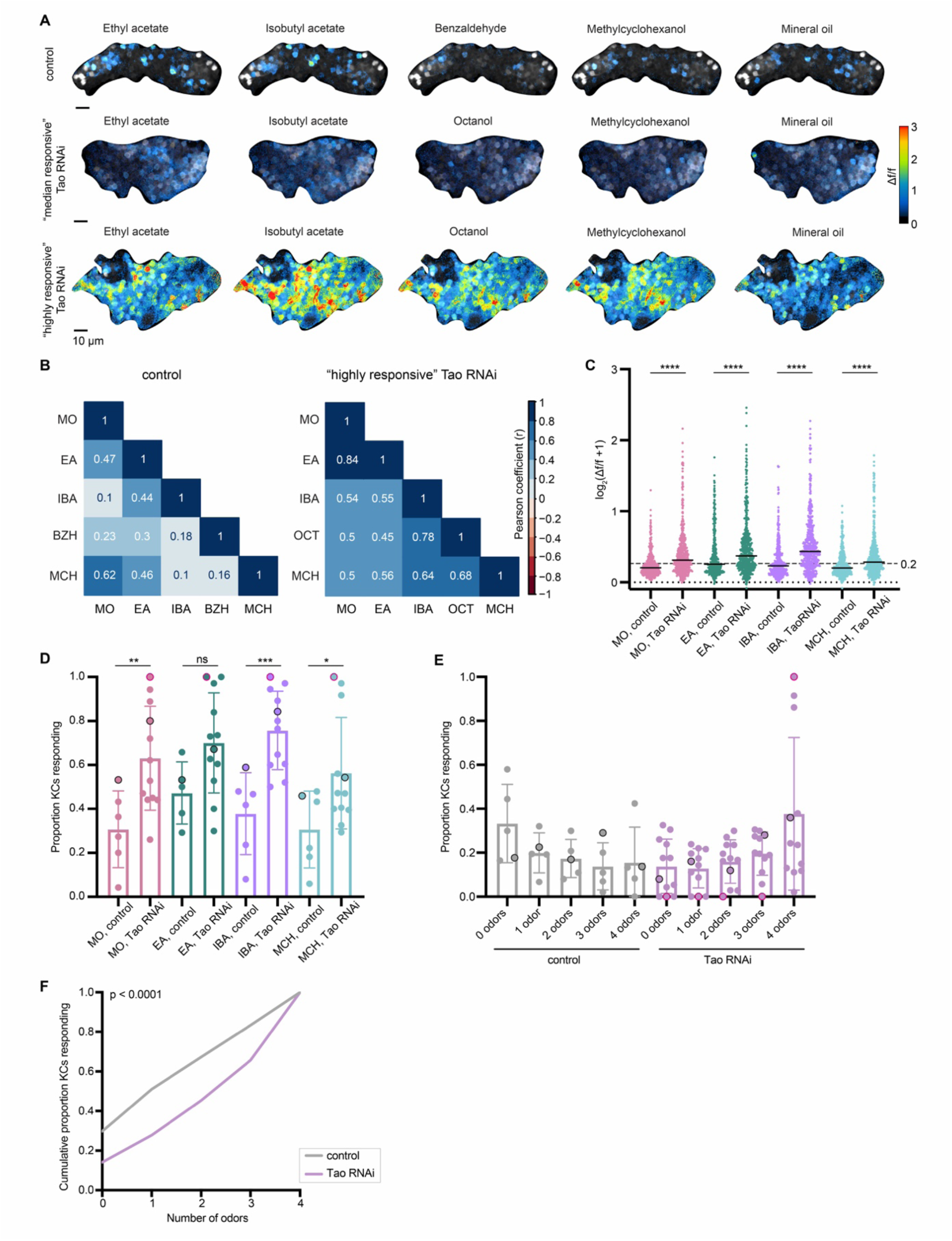
Odor selectivity is reduced in Kenyon cells with increased dendritic claw numbers. (A) Example KC somatic odor responses in control hemisphere and KC>*Tao* RNAi (increased dendrite condition) hemispheres with median levels of odor responses, and high levels of odor responses. Grayscale backdrop indicates the cells. Δf/f scale is shown such that all cells with responses < 0.2 are colored black, responses > 3 are colored red. Control dataset shown throughout this figure are the same data as controls for *mud* RNAi condition in Figure 1, and the sample shown in (A) is the same sample shown in Figure 1G. In *Tao* RNAi animals, benzaldehyde was switched with octanol due to a crystallization issue with benzaldehyde in the olfactometer. Further discussion is provided in the methods. See also Figure S3 for odor response traces over time for the control and “highly responsive” *Tao* RNAi images. (B) Pearson correlation matrix (half diagonal displayed) show the linear relation between each pairwise odor comparison across the cells for the control and “highly responsive” KC>*Tao* RNAi image shown in A. Correlation value (r) can fall in a range of –1 (dark red) to +1 (dark blue). However, we did not observe any negative correlations in this analysis. (C) Peak odor responses of all cells, aggregated from all analyzed samples. Y axis displays log_2_-scaled Δf/f values. Dashed line indicates 0.2 Δf/f threshold. Dotted line indicates Δf/f = 0. Black horizontal bars show medians. Significance: Mann-Whitney test. n=349-404 control cells, 496 *Tao* RNAi cells for each odor. (D) Proportion of cells in each sample responding to each odor above 0.2 Δf/f threshold. Significance: unpaired t-test. (E) For samples in which the same cells could be tracked across all odor presentations, proportion of cells responding to 0, 1, or multiple odors is shown. Bar plots in (D,E) show mean ± SD, and black circled data points correspond to the control, and “median responsive” *Tao* RNAi images shown in (A). Red circled data points correspond to the “highly responsive” *Tao* RNAi image in (A). (F) Cumulative proportion of cells responding from 0 up to 4 odors. Lines represent mean of all control (gray; n = 349) and *Tao* RNAi cells (purple; n = 496). Significance: Kruskal-Wallis test; K-S distance = 0.2318.

We next asked what proportion of Kenyon cells per calyx responded to 0, 1 or multiple odors. We observed a sharp increase in cells that responded to all odors, from <20% in controls to a mean of 40% in increased-dendrite animals (Figures 3E, F, S3B). Taken together, we see a striking change in sensory coding: odors no longer activate sparse populations of Kenyon cells, and individual cells are no longer odor selective. In previous work, a similar change in olfactory coding was induced by acute inhibition of the GABAergic interneuron APL, which scales population-level odor responses to overall sensory drive (Lin et al., 2014). Loss of APL feedback prevented animals from discriminating among odors. Our manipulation is the first hard-wired alteration to Kenyon cell input number and demonstrates the significance of input number in tuning sensory selectivity of expansion layer neurons.

### Overexpressing dendritically-targeted Dscam1 in Kenyon cells decreases dendritic claws per cell

We next asked if we could manipulate Kenyon cell dendrites in the opposite direction, reducing claw number per cell. Very few genes have been shown to traffic specifically to dendrites or to affect only dendrite structure without affecting axon structure. One such gene is the dendritically targeted isoform of Dscam1: While Dscam1 isoforms containing transmembrane domain 2 (TM2) traffic to both axons and dendrites, Dscam1-TM1 isoforms preferentially traffic to dendrites (Wang et al., 2004) . Ectopic expression of single TM1 isoforms in Kenyon cells was previously shown to disrupt calyx morphology without altering the morphology of mushroom body lobes (Wang et al., 2004) . Therefore, we asked if the changes in calyx morphology are a result of an effect on dendritic arborization of KCs. We will refer to the GFP-tagged Dscam1[TM1] variant as Dscam^3.36.25.1^ (Wang et al., 2004). We overexpressed Dscam^3.36.25.1^ in Kenyon cells and sparsely labeled them using mBitbow 2.2 to visualize individual dendritic claws (Figure 4A). There was a drastic decrease in the number of claws per Kenyon cell, from a median of 7 in controls to 1 in Dscam^3.36.25.1^ animals (Figure 4B). While Kenyon cell number did not change significantly, the calyx area became significantly smaller (Figure 4C-F).

**Figure 4.**
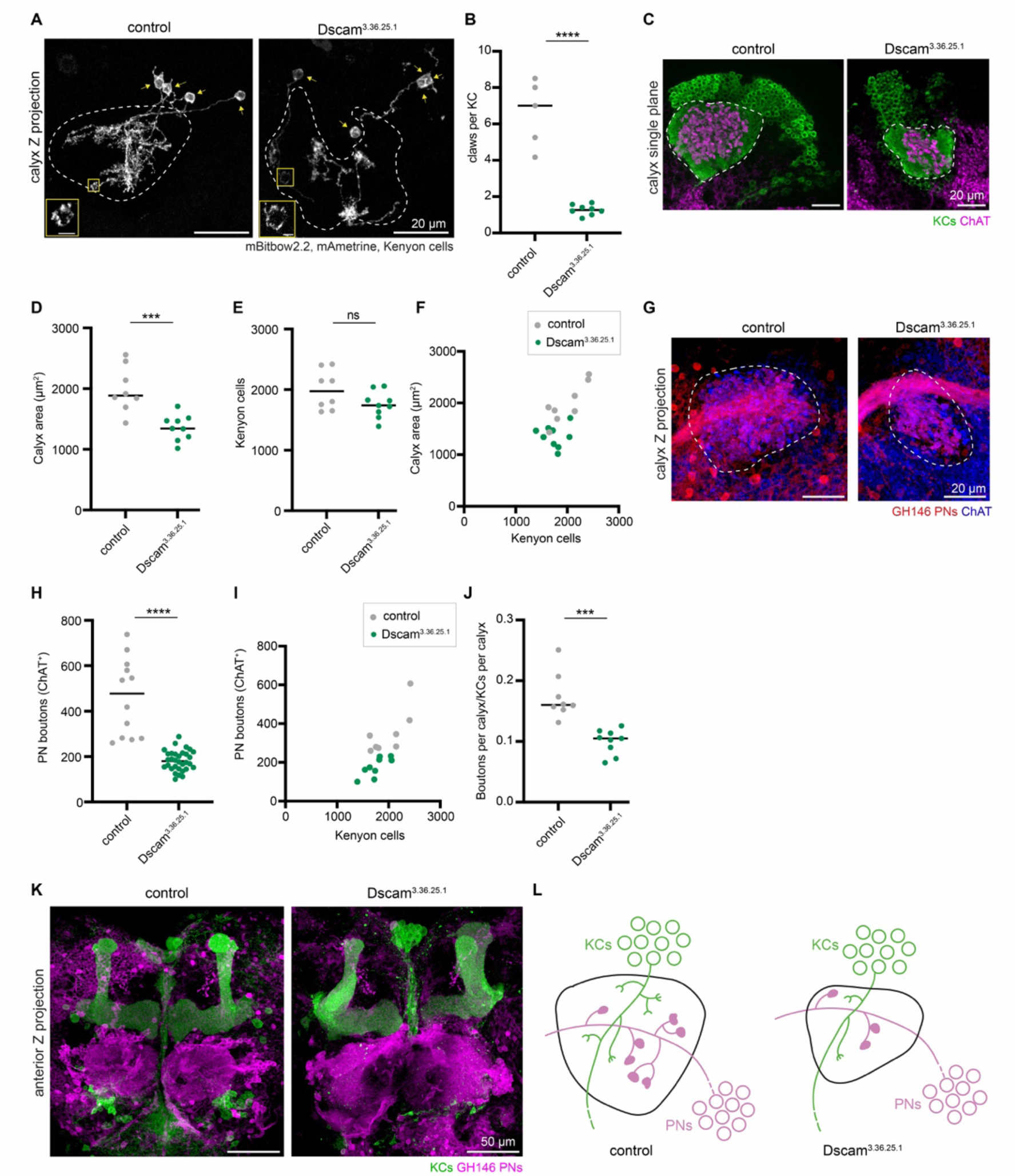
Overexpressing dendritically-targeted Dscam1 in Kenyon cells decreases dendritic claws per cell. (A) Example maximum intensity Z-projections of sparsely labeled KCs in control and KC>*Dscam^3.36.25.1^* (Dscam1[TM1]) calyces. Dashed lines outline the calyx, and arrows mark the KC soma. KCs labeled by mBitbow2.2 (mAmetrine shown). Insets zoom in on claw structure from the yellow boxed region in the corresponding image, with brightness increased for visualization. Scale bar for inset: 2μm. (B) Number of dendritic claws per KC in control (gray) and KC>*Dscam^3.36.25.1^* (green) hemispheres. (C) Confocal slice of control calyx and KC>*Dscam^3.36.25.1^* calyx. OK107 labels KCs (green) and ChAT labels projection neuron boutons (magenta). Representative images selected with median number of boutons from each condition, and image slice chosen to show a middle plane of boutons. Dashed lines outline the calyx. (D-F) Maximum calyx cross-sectional area (D), number of KCs (E), and the relationship between KC number and maximum calyx cross-sectional area (F) in control (gray) and KC>*Dscam^3.36.25.1^* hemispheres (green). (G) Maximum intensity projection of confocal stack of calyx bouton production by GH146^+^ PNs (red) in control and KC>*Dscam^3.36.25.1^* hemispheres. ChAT immunostaining (blue) highlights calyx extent (circled). (H-J) PN bouton number (H,I), and ratio of ChAT^+^ boutons per KC (J) in control (gray) and KC>*Dscam^3.36.25.1^* hemispheres (green). (K) Maximum intensity projection of confocal stacks taken from anterior side of control and KC>*Dscam^3.36.25.1^* brains. KCs (green) show the mushroom body lobes, GH146^+^ PNs (magenta) show the antennal lobe and PN cell bodies. (L) Model of the effect of decreasing KC dendritic claws per cell. PN bouton production decreases, and the PN tract shifts dorsally. Throughout this figure, significance: unpaired t-test, and black horizontal bars indicate the median.

To ask how the reduction of Kenyon cell dendritic claws affects PN bouton numbers, we quantified ChAT-labeled boutons (Figure 4G, H). Median bouton numbers in Dscam^3.36.25.1^ animals declined by ∼½ (Figure 4H-J). Similar to the increased-dendrite condition, we observed a shift of the PN axon tract in the Dscam^3.36.25.1^ calyces, this time in the dorsal direction which could suggest a change in spatial distribution of PN boutons in the calyx (Figure 4G). KC axons and antennal lobes retained normal gross morphology (Figure 4K). Overall, these anatomic effects were reciprocal to those in the Tao knockdown condition (Figures 2O, 4L); we therefore tested if odor responses changed reciprocally as well.

### Kenyon cells with reduced claw numbers are less responsive to odors

Dscam^3.36.25.1^ Kenyon cells, with reduced dendrites, displayed reduced odor responses compared to controls: About half as many cells responded to each odor (Figures 5A-C, S5A). Next, we asked what proportion of Kenyon cells per calyx responded to 0, 1 or multiple odors. We observed a sharp increase in cells that responded to no odors, from ∼20% in controls to ∼60% in reduced-claw calyces (Figures 5D-E, S5B).

**Figure 5.**
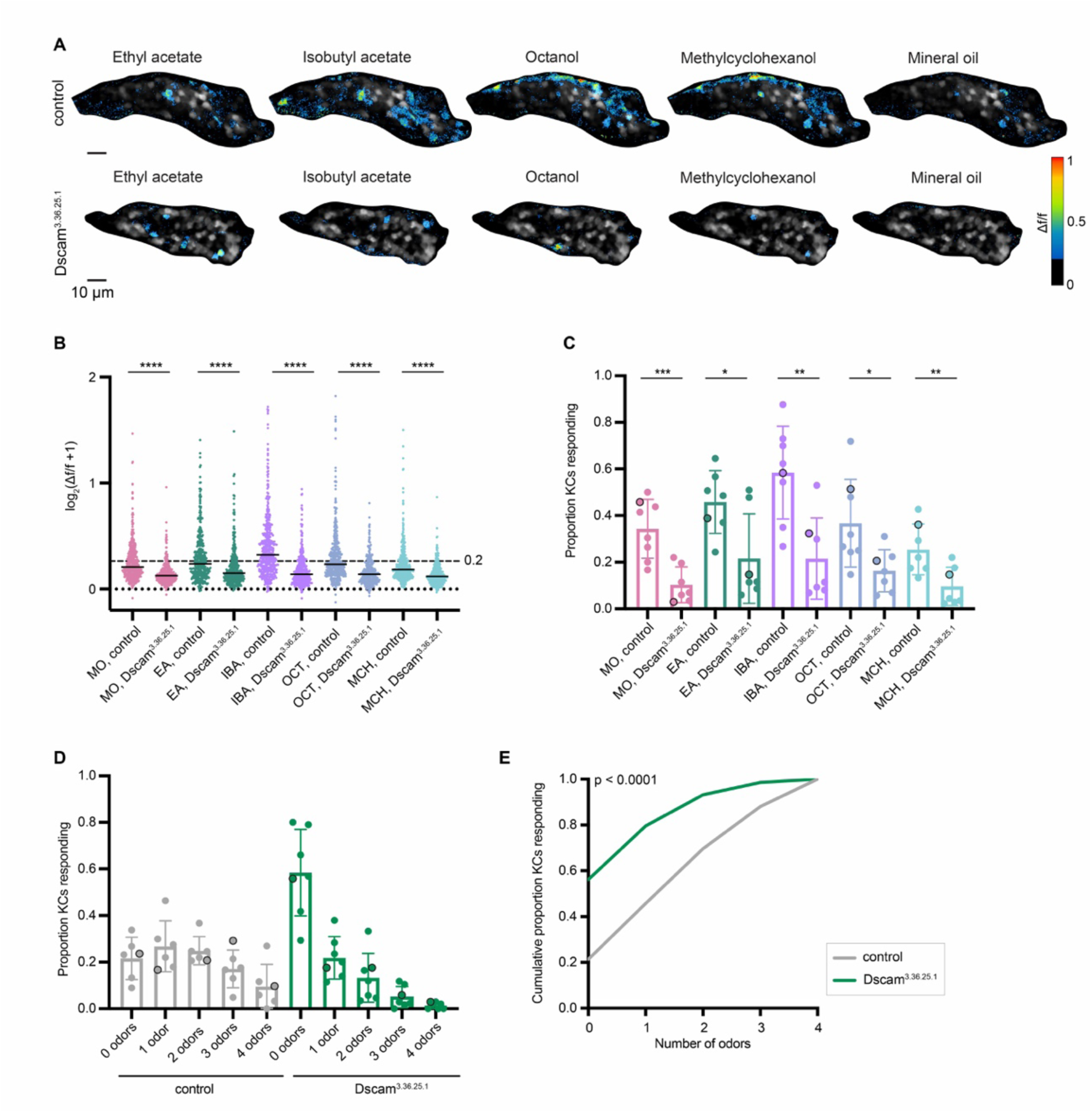
Kenyon cells with reduced claw numbers are less responsive to odors. (A) Example KC somatic odor responses in control hemisphere and KC>*Dscam^3.36.25.1^* (decreased dendrite condition) hemisphere. Grayscale indicates the cells. Δf/f scale is shown such that all cells with responses < 0.2 are colored black, responses > 1 are colored red. See also Figure S5 for odor response traces over time for this image. (B) Peak odor responses of all cells, aggregated from all analyzed samples. Y axis displays log_2_-scaled Δf/f values. Dashed line indicates 0.2 Δf/f threshold. Dotted line indicates Δf/f = 0. Black horizontal bars show medians. Significance: Mann-Whitney test. n = 380-428 control cells, 372 Dscam^3.36.25.1^ cells for each odor. (C) Proportion of cells in each sample responding to each odor above 0.2 Δf/f threshold. Significance: unpaired t-test. (D) For samples in which the same cells could be tracked across all odor presentations, fraction of cells responding to 0, 1, or multiple odors is shown. Bar plots in (C,D) show mean ± SD, and black circled data points correspond to the images shown in (A). (E) Cumulative proportion of cells responding from 0 up to 4 odors. Lines represent mean of all control (gray; n = 369) and Dscam^3.36.25.1^ cells (green; n = 338). Significance: Kruskal-Wallis test; K-S distance = 0.348.

Altogether, we see a striking change in odor coding in the decreased-dendrite animals: Kenyon cells become less responsive to odors, leading to population-level Kenyon cell responses becoming sparser than controls. This demonstrates that decreasing the number of dendritic claws per Kenyon cell and hence the number of odor inputs makes Kenyon cell odor responses sparser and more selective.

### Animals with reduced numbers of Kenyon cells exhibit associative learning in a Y-arena two-odor choice assay

To allow association of odor information with contingent events, Kenyon cells connect with dopaminergic neurons (DANs) and mushroom body output neurons (MBONs) in the mushroom body lobes. DANs convey environmental context and can signal positive and negative events; coincident firing of Kenyon cells and DANs results in alteration of Kenyon cell-MBON synaptic weight and modulates future behavioral responses to the same odor stimulus (Aso and Rubin, 2016; Cohn et al., 2015; Handler et al., 2019; Hige et al., 2015; Owald and Waddell, 2015; Séjourné et al., 2011; Tomchik and Davis, 2009). We have shown that Kenyon cell input number and sparse odor coding are preserved in mushroom bodies composed of 500-4000 Kenyon cells per hemisphere ((Elkahlah et al., 2020) and Figure 1 above). We therefore sought to ask how the number of Kenyon cells in the brain influences associative learning abilities.

In *D. melanogaster*, four mushroom body neuroblasts from each hemisphere give rise to ∼500 KCs each (Ito et al., 1997). To generate flies with reduced Kenyon cell numbers, we use hydroxyurea to ablate Kenyon cell neuroblasts shortly after larval hatching (de Belle and Heisenberg, 1994; Elkahlah et al., 2020; Sweeney et al., 2012). Previous work has shown that groups of animals treated with high hydroxyurea dose lose all 8 neuroblasts and cannot form odor associations at all (de Belle and Heisenberg, 1994). Mild hydroxyurea treatment results in stochastic Kenyon cell neuroblast loss such that individual treated animals have between 0 and 8 Kenyon cell clones between the two hemispheres ((Elkahlah et al., 2020) and our results here). Groups of animals with mild hydroxyurea treatment have degraded associative learning performance (de Belle and Heisenberg, 1994). We therefore sought to relate the associative learning abilities and Kenyon cell numbers of *individual* flies.

To do this, we tested individual hydroxyurea-treated flies in a recently described two-alternative forced choice task (2AFC) (Figure 6A) (Rajagopalan et al., 2022). This task utilizes a Y-shaped arena, in which we measured a fly’s choices of two different odors, 3-octanol and 4-methylcyclohexanol. Each fly was dissected post-hoc and immunostained to score the number of Kenyon cell neuroblast clones (Figures 6B, S6A). To facilitate the counting, we labeled the latest born ‘αβ core’ KCs. The somata and neurites of these Kenyon cells form four spatially separable clusters, allowing us to score Kenyon cell clone number by counting the groups of labeled soma or axon tracts (Elkahlah et al., 2020). As hydroxyurea ablation sometimes affects the lNB/BAlc that gives rise to lateral PNs, we included the MZ19 marker to track the lateral DA1 glomerulus (Lai et al., 2008; Stocker et al., 1997). Almost all animals retained at least one PN neuroblast (Figures S6C, D), while the number of Kenyon cell clonal units varied widely.

**Figure 6.**
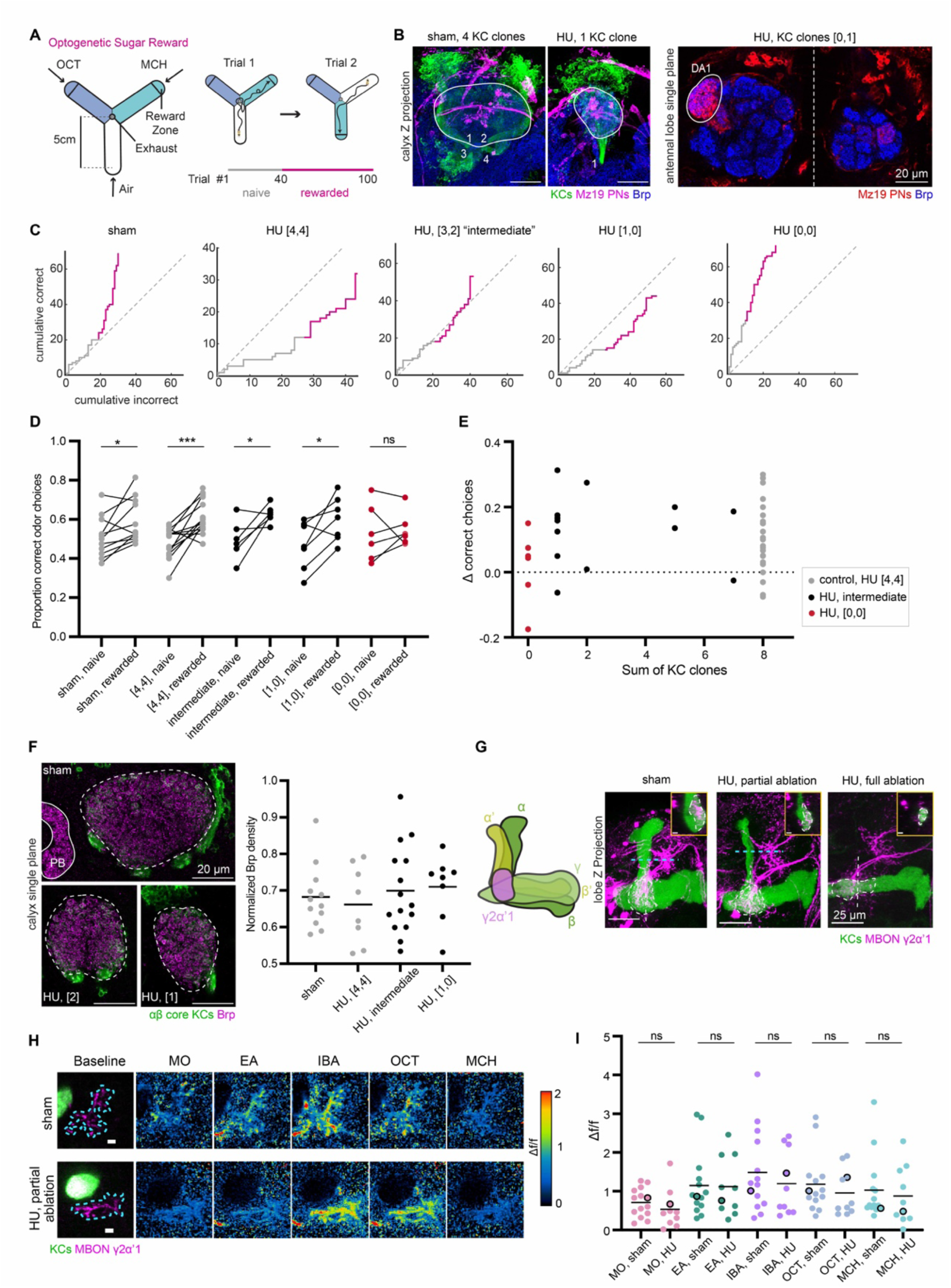
Animals with reduced numbers of Kenyon cells exhibit normal behavioral and feed-forward functional responses. (A) Schematic of Y-arena for single fly learning experiments (left). Airflow travels from tips of each arm to an outlet in the center. Reward zones are indicated by lines on the schematic, but are not visible to the fly. A choice is considered to have been made when a fly crosses from an air arm into the reward zone of an odorized arm, triggering delivery of an optogenetic reward with a 500 ms pulse of red light. The next trial then commences as the choice arm switches to air and the two odors (represented with two different colors for octanol and methylcyclohexanol) are randomly reassigned to the other two arms (right). Naïve (1-40) and rewarded trials (41-100) are indicated below. (B) Left: Maximum intensity projections of confocal stacks of calyces of sham-treated and HU-treated animals. 58F02 labels αβ core KCs (green). Mz19 labels a subset of ∼16 PNs (magenta). Brp (blue) is included as a neuropil marker. Calyx is outlined in white. The 58F02 signal allows scoring of the number of KC clones. Numbers (1 through 4 in sham, 1 in HU-treated calyx) are shown to illustrate the number of neurite bundles innervating the pedunculus when traced through the stack. Right: Single confocal slice of the left and right antennal lobe from a HU-treated brain that has 1 KC clone in the left hemisphere, and no clones on the right side. Dotted line indicates the midline of the central brain. Mz19 labels the DA1 glomerulus (encircled in white), allowing scoring of the lNB/BAlc PN neuroblast. The lNB/BAlc PN neuroblast is ablated in the right hemisphere and DA1 glomerulus is lost. (C) Learning curves (gray lines) of individual flies from each condition are plotted to show cumulative correct (rewarded) odor choices and incorrect (unrewarded) odor choices across all 100 trials. Some flies were tested with a different number of trials, e.g. the HU, “[4,4]” fly shown here; details of the behavior protocol are explained in the Methods section. Naïve (gray line) and rewarded (pink line) trials are indicated. Gray dotted line displays y=x line through the origin, indicating equal preference for the two odors. (D) Proportion of correct odor choices made in naïve and rewarded trials; learning is indicated if the number of trials with correct odor choices is higher in the rewarded trials than in naïve trials. Each data point is an individual fly. Sham-treated animals, and HU-treated animals with “[4,4]” KC clones are displayed in gray, while intermediate HU-treated clone numbers are shown in black, and fully ablated “[0,0]” HU-treated animals are shown in red. Jitter added in this plot and (E) to display all the data points. Significance: paired t-test. See also Figure S6B. (E) Relationship between Δ correct choices (difference in correct choices made in rewarded vs naïve trails) and sum of KC clones from both hemispheres (range of 0-8). No correlation was observed. See Figure S6C-D for relation with lNB/BAlc ablation. (F) Left: Confocal slice with maximum calyx cross-sectional area of sham and HU-treated calyces (bottom left: 2 KC clones, bottom right: 1 KC clone). Dashed lines outline the calyx. Part of the normalization structure, protocerebral bridge (PB) is included in the sham image. Right: Quantification of Bruchpilot density, normalized to fluorescence in PB. Black bars represent medians. All pairwise comparisons are non-significant by unpaired t-test. See also Figure S6E. (G) Left: Schematic of the mushroom body lobe anatomy with KCs in green and MBONs γ2α’1 in magenta. The different shades of green represent distinct MB lobes named due to KC axonal morphology differences. Dendrites of MBONs γ2α’1 in the lobe compartment are shown. Right: Maximum intensity projections of confocal stack of MB lobe with KCs (green) and γ2α’1 MBON (magenta) labeled in sham; HU-treated: partially ablated; and HU-treated: fully ablated animals. In each image, the MBON compartment is circled with a dashed line. Inset shows the lateral view of the MBON compartment. Scale bar for inset: 5 µm. Corresponding location is indicated by white dashed vertical line. Blue dashed line indicates the focal plane for calcium imaging of MBON axonal terminals in (H, I). See also Figure S6A for effect on arborization of PAM-DANs in the mushroom body lobes. (H) Example γ2α’1 MBON axonal odor responses in sham hemisphere and HU-treated: partially ablated hemisphere. Left: Focal plane of KC vertical lobe (green) and resting GCaMP6s signal in MBON axonal terminals (magenta). Blue dashes outline the ROI used to measure odor responses. Scale bar: 5 µm. Right: Responses to odors in Δf/f. Δf/f scale is shown such that all pixels with responses < 0 are colored black, responses > 2 are colored red. (I) Peak odor responses of all MBON axonal ROIs. Each dot is one hemisphere. For HU-treated: partially ablated hemispheres, only hemispheres with maximum cross-sectional calyx area less than every control (< 2100 µm^2^) are included (Figure S6G). Black horizontal bars show mean peak odor responses. Significance: unpaired t-test. Black circled data points correspond to samples shown in (H). Average traces of odor responses can be found in Figure S6F.

Our behavioral apparatus allowed us to measure odor choices of individual flies over many trials. Each behavior session is initialized so the Y-arm containing the fly is filled with air and odors randomly assigned to the other two arms. When the fly travels to the end of one of the odorized arms, this is defined as a choice, and reward was provided by optogenetically activating reward signaling protocerebral anterior medial (PAM)-DANs (Burke et al., 2012; Liu et al., 2012). The arena then resets, the arm chosen by the fly switches to clean air, the other two arms are randomly assigned the two odors and the cycle of trials continues. For each session, we first assessed the naïve odor choices of each fly for either 40 or 60 trials, before measuring learning by reinforcing one of the odors, chosen at random for each experiment, over 40 or 60 subsequent trials. This allowed us to account for any bias in the choices of an individual animal by calculating the change in choice probability before and after reward is available.

With PAM-DAN reinforcement, sham-treated control animals chose the rewarded odor in 59% of trials on average, compared to 51% prior to training (Figures 6D, S6B) (Rajagopalan et al., 2022). Hydroxyurea-treated animals that retained all four Kenyon cell clones in both hemispheres (“[4,4]” set) learned similarly to controls, while animals with all 8 clones lost (”[0,0]”) failed to learn (Figures 6D, S6B). We then evaluated the effects of intermediate reductions in Kenyon cell number. To our surprise, flies with at least one remaining Kenyon cell clone (“[1,0]”) were still able to increase their preference for the odor paired with PAM-DAN reward. As a result, learning performance did not correlate with number of Kenyon cell clones (Figure 6E). This is consistent with results that indicate that as few as 25 Kenyon cells can be adequate for computationally distinguishing different odors, as well as behavioral effects of partial mushroom body loss in honeybees (Campbell et al., 2013; Malun et al., 2002).

Brp staining of these behaviorally-tested flies indicated that synaptic density remained consistent across animals with different numbers of Kenyon cells (Figures 6F, S6E). This suggests the number of synapses per Kenyon cell are not altered in these partially-ablated flies, and is consistent with our previous findings that population-level odor responses in these animals are similar to controls (Elkahlah et al., 2020). We note that the odor choices we used for this task were very distinct, and it is possible that animals with reduced Kenyon cell repertoires would fail to discriminate chemically similar odors or mixtures. However, since PAM-DAN optogenetic reinforcement produced relatively weak learning scores in control animals, we did not attempt to subject Kenyon cell-ablated animals to more difficult discrimination tasks.

### Anatomic and functional properties of downstream circuitry in flies with reduced Kenyon cell numbers

The ability of an animal to associate an odor with a reward depends on tripartite synaptic connections between Kenyon cells, MBONs, and DANs in the mushroom body lobes. As animals with reduced Kenyon cell number retain this behavioral ability, we hypothesized that the connections of Kenyon cells to MBONs and DANs is intact in these animals. The axon bundles of KCs form “L”-shaped mushroom body lobes (Figure 6G), including vertical α and α’ lobes and horizontal β, β’, and γ lobes. Based on dendritic arborization of MBONs and axonal projection patterns of DANs, the mushroom body lobes can be further divided into 15 different compartments (Aso et al., 2014a). Within a certain compartment, individual MBONs receive inputs from Kenyon cells, while compartment-specific DANs modulate connection strength between KCs and MBONs (Handler et al., 2019; Li et al., 2020). These precise compartmental divisions are expected to allow precise behavioral responses to conditioned stimuli (Aso et al., 2014b; Eschbach et al., 2020; Scaplen et al., 2021).

To evaluate the potential for this circuit to support learning with these reduced Kenyon cell numbers, we examined the anatomy and functional response properties of downstream circuitry required for learning. The Mz19 driver used to assess the specificity of our HU ablations also labels PAM-DANs that project their axons to the β’2γ5 compartment of the mushroom body (Aso et al., 2012). This fortuitous situation allowed us to assess whether DAN innervation of the mushroom body was affected when Kenyon cell number was reduced through neuroblast ablation (Figure S6A). In flies with reduced Kenyon cell numbers, β’2γ5 PAM-DANs project to the same region as controls. Dopaminergic projections were even preserved in fully ablated flies that only retained embryonically-born γd KCs. Overall, the anatomy was compatible with the possibility that learning/dopaminergic teaching signals might still be intact after these manipulations.

To examine MBON anatomy and function, we chose the γ2,α’1 MBON due to its genetic and anatomic compatibility with all our Kenyon cell manipulations (described further below). The activation of γ2,α’1 MBON is sufficient to drive approach behavior, in line with our reward learning behavior assay (Aso et al., 2014b). To examine the functional connections between Kenyon cells and MBONs after manipulations, we used R25D01-LexA to express GCaMP6s in γ2,α’1 MBON (Figure 6G). Dendrites of γ2,α’1 MBON arborize at the intersection of vertical and horizontal lobes of the mushroom body, and receive inputs from both γ and α’ lobes (Li et al., 2020).

We first visualized the morphology of γ2,α’1 MBON in hydroxyurea-treated reduced-Kenyon cell animals. Despite the shrunken lobes in these animals, the γ2,α’1 MBON still arborized at the intersection of vertical and horizontal lobes. γ2,α’1 MBONs colocalized with KCs in the γ lobe similar to shams (Figure 6G, see inset). In fully ablated-Kenyon cell animals, only embryonic-born γd KCs are retained to form part of the γ lobe (Armstrong et al., 1998). γ2,α’1 MBON dendrites were seen throughout the γ lobe in the transverse plane and slightly extended along the lobe in the coronal z-projection (Figure 6G). Consistent with a recent study of MBON morphology in HU-fully ablated animals, we also observed γ2,α’1 MBON dendrites to arborize more sparsely in the lobes, with slight extension outside the lobes (Lin et al., 2022).

Given the normal dendritic arborization of γ2,α’1 MBON in partially ablated animals, we asked whether these KC-MBON connections are functional. To do this, we imaged calcium activity of γ2,α’1 MBON axonal arbor in response to different odors and mineral oil (Figure 6H). We measured calcium responses in axon terminals so as to measure MBON activity that is transmitted to downstream neurons to modulate behaviors. For all the odors tested, peak odor responses were comparable between sham and partially ablated animals (Figures 6H, I, S6F).

Although calyx area predicts Kenyon cell number in this condition, we did not observe any correlation between peak odor response and the calyx area (Figure S6G) (Elkahlah et al., 2020). This indicates that even a reduced population of KCs can successfully transmit signals to the γ2,α’1 MBONs.

### Associative learning abilities of animals with increased Kenyon cell number or claw number

We next tested the learning capabilities of individual OK107> *mud* RNAi and *Tao* RNAi flies, with expanded Kenyon cell repertoire or claw number, respectively. Again, the learning scores of individual flies were assessed and their mushroom body anatomy evaluated by post-hoc staining. In this set of experiments, we provided optogenetic reward using the Gr64f population of sugar-sensing gustatory neurons, which produced much better learning scores in control animals than PAM-DAN reinforcement (Figure 7A); an explanation for this difference is discussed in (Rajagopalan et al., 2022). The proportion of trials with correct odor choices was quantified in naïve and rewarded trials (Figure 7B, C). All three genotypes showed robust, odor-specific learning. Because Kenyon cell augmentation was highly variable in the *mud* RNAi genotype, we measured the relationship between calyx size and learning score. Surprisingly, learning improved with increasing calyx size, suggesting that increasing the size of the Kenyon cell repertoire gives animals more power to discriminate odors, form odor-specific learned associations, or use learned associations to guide decisions (Figure 7D). The robust odor associations formed by flies with excess Kenyon cell dendrites is also surprising, as the weakened Kenyon cell odor selectivity in these animals would theoretically make odor discrimination more difficult. It is possible that this behavioral task is relatively simple, and these flies may fail to learn in a more difficult task of discriminating molecularly similar odors or mixtures.

**Figure 7.**
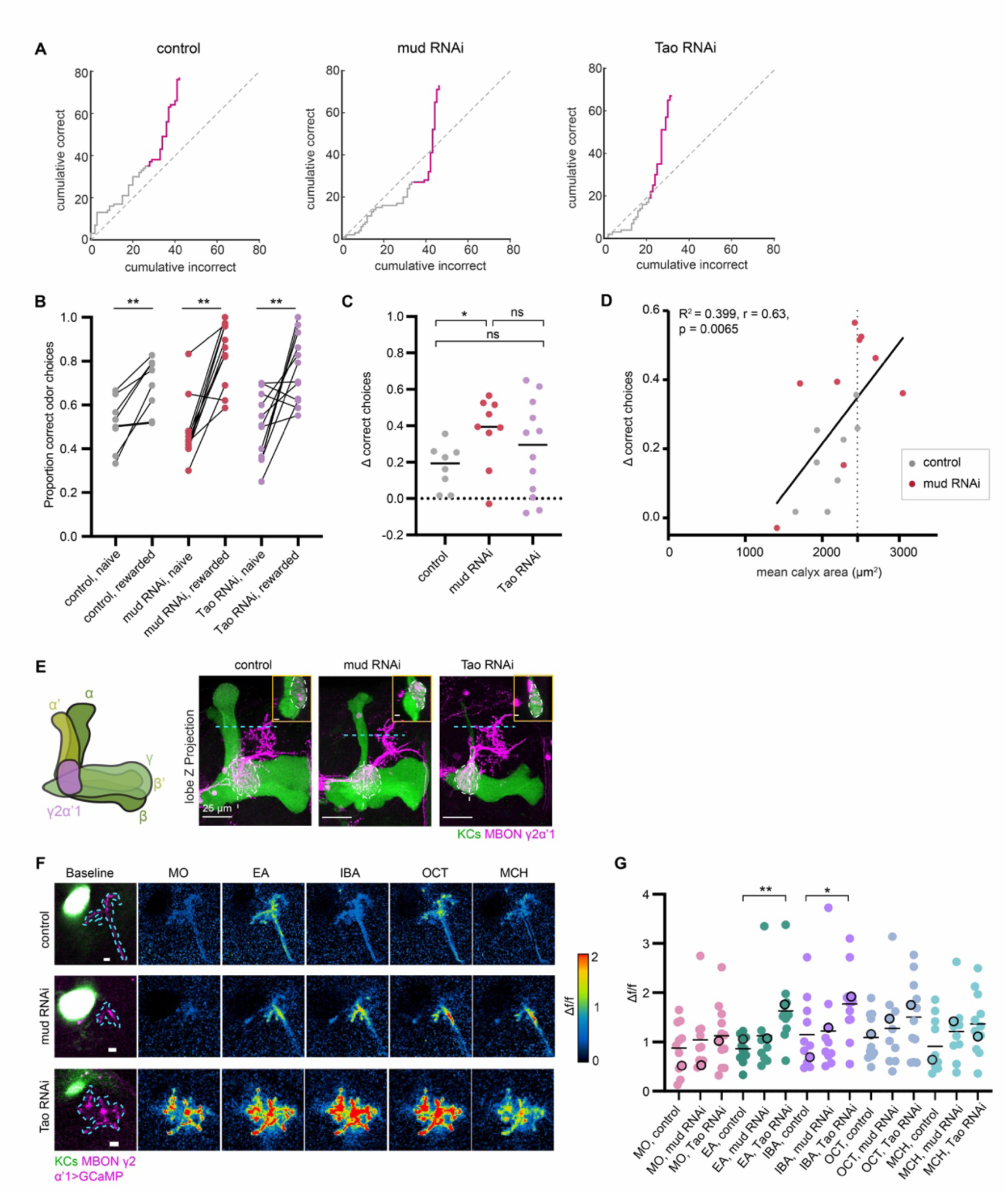
Effect of increased Kenyon cell number or claw number on associative learning and feedforward functional responses. (A) Learning curves (gray lines) of individual flies from control, KC>*mud* RNAi, and KC>*Tao* RNAi are plotted to show cumulative correct and incorrect odor choices across all trials. Naïve (gray line) and rewarded (pink line) trials are indicated. Flies were either tested with 40 or 60 naïve and rewarded trials each (details explained in Methods). Gray dotted line displays y=x line through the origin. (B) Proportion of correct odor choices made in naïve and rewarded trials, in control (gray), KC>*mud* RNAi (red), and KC>*Tao* RNAi (purple) animals. Each data point is an individual fly. Jitter added in this plot and (C) to display all the data points. Significance: paired t-test. (C) Δ correct choices (difference in correct choices made in rewarded vs naïve trails) in control (gray), KC>*mud* RNAi (red), and KC>*Tao* RNAi (purple) animals. Dotted line at y = 0 indicates no change in odor preferences. Significance: unpaired t-test. (D) Δ correct choices plotted against mean calyx area for each animal in control and KC>*mud* RNAi conditions. Black line represents linear fitted line. Dotted line represents the largest control calyx. (E) Maximum intensity projections of confocal stack of MB lobe with KCs (green) and MBONs γ2α’1 (magenta) labeled in control, *mud* RNAi and *Tao* RNAi animals. In each image, the MBON compartment is circled with a dashed line. Inset shows the lateral view of the MBON compartment at the location of the white dashed vertical line (inset scale bar: 5 µm). Blue dashed line indicates the focal plane for calcium imaging of MBON axonal terminals in (F, G). (F) Example MBON axonal odor responses in control, *mud* RNAi and *Tao* RNAi animals. Left: Focal plane of KC vertical lobe (green) and resting GCaMP6s signal in MBON axonal terminals (magenta). Blue dashes outline the ROI used to measure odor responses. Scale bar: 5 µm. Right: Responses to odors in Δf/f. Δf/f scale is shown such that all pixels with responses < 0 are colored black, responses > 2 are colored red. (G) Peak odor responses of all MBON axonal ROIs for the three conditions; each dot is one hemisphere. For *mud* RNAi hemispheres, only hemispheres greater than every control (maximum cross-sectional calyx area > 2200 µm^2^) are included (Figure S7B). Black horizontal bars show mean peak responses. Significance: Peak odor responses of *mud* RNAi and *Tao* RNAi are compared with control using unpaired t-tests. Non-significant comparisons are not shown. Black circled data points correspond to responses from the images in (F). Average traces of odor responses can be found in Figure S7A.

### MBON odor responses are robust to an increase in Kenyon cell number and Kenyon cell claw number

Next, using the same method, we examined functional connections between Kenyon cells and γ2,α’1 MBON in the increased-Kenyon cell and increased-Kenyon cell claw animals (OK107> *mud* RNAi, OK107>*Tao* RNAi, and OK107>empty attp2 RNAi control). We chose the γ2,α’1 MBON because the γ lobe is the main mushroom body lobe retained in *Tao* RNAi animals (Figure 2N) (King et al., 2011). On rare occasions, the vertical mushroom body lobes were also shrunken in *mud* RNAi animals due to mislocalization of axons to the calyx. In our experiments, genetic background strongly influences the rate of axon mislocalization in *mud* RNAi knockdown (Figure 7E). Dendrites of the γ2,α’1 MBON projected to the same compartment in the *mud* RNAi and *Tao* RNAi animals as in controls (Figure 7E). In OK107>*Tao* RNAi animals, γ2,α’1 MBON dendrites were seen throughout the remaining γ lobe (Figure 7E, inset). We next measured odor responses in the γ2,α’1 MBON axon terminal in both *mud* RNAi and *Tao* RNAi animals. Interestingly, the MBONs of Kenyon cell *mud* RNAi and *Tao* RNAi animals responded to all four odors tested (Figure 7F). Peak odor responses to the same odor were comparable between control and *mud* RNAi animals (Figures 7G, S7A). Strikingly, for two of the odors tested (EA and IBA), peak odor responses were significantly higher in *Tao* RNAi animals compared to control animals (Figure 7G, S7A), consistent with our result above that Kenyon cells of *Tao* RNAi animals receive increased inputs from PNs (Figure 3). We did not observe a correlation between MBON odor responses and calyx area in OK107> *mud* RNAi animals (Figure S7B).

In conclusion, γ2,α’1 MBON shows robust functional input from Kenyon cells regardless of manipulation of Kenyon cell number or claw number. This provides an anatomical basis for the robust learning behavior we reported in these conditions.

## Discussion

Here, we have initiated a new method, developmental hacking of circuit wiring, to test how specific circuit parameters of the arthropod expansion layer influence cognitive computations. Using chemical and genetic approaches, we increase and decrease Kenyon cell number and Kenyon cell input number. This approach adds to a growing body of literature in which researchers engineer cell biological changes to neurons in order to probe developmental algorithms or change circuit function (Dzirasa et al., 2022; Hawk et al., 2021; Heckman and Doe, 2022; Linneweber et al., 2020; Meng et al., 2019; Pechuk et al., 2022; Pop et al., 2020; Prieto-Godino et al., 2020). Both the developmental and functional results force us to rethink assumptions and predictions about how the structure of connectivity influences odor representations and learned associations. First, we find that changing Kenyon cell number, and thus expansion ratio, has minimal effect on Kenyon cell odor representation. Moreover, animals with diminished Kenyon cell repertoires can still make simple learned associations, and animals with larger numbers of Kenyon cells show improved two-choice odor learning. These surprising findings suggest that the developmental algorithms we and others identified previously, which prioritize relationships among circuit layers rather than cellular precision, allow nervous systems to be computationally robust (Elkahlah et al., 2020; Kiral et al., 2021; Otopalik et al., 2017). Second, we find that lowering Kenyon cell input density makes Kenyon cells more stringent in their odor responses, while raising input density makes them more promiscuous. Again, this result is informative about the developmental relationship between input density and physiological response threshold. Shockingly, animals with *Tao* knocked down in Kenyon cells, which results in excess Kenyon cell claws, promiscuous Kenyon cell odor responses, and disrupted Kenyon cell axons, remain adept at a two-choice odor learning task.

As our understanding of mushroom body development becomes more sophisticated, we will illuminate *how* these computational variables and cellular relationships are specified and gain increasingly nuanced control over their *in vivo* implementation.

### Characteristics of expansion layer neurons govern both input and output weights

While neurons of all expansion layers receive broad sensory inputs, Kenyon cells of the mushroom body have the special property of connecting all local sensory inputs to a multitude of neurons that govern behavioral outputs. This keystone position in the circuit gives them an unusual developmental relationship to the pre- and post-synaptic cells with whom they synapse. In previous work, we showed that the connectivity relationship between presynaptic olfactory PNs and postsynaptic Kenyon cells is set by the Kenyon cells. When we varied Kenyon cell number and PN number, the number of inputs to individual Kenyon cells always averaged 5-6 per cell (Elkahlah et al., 2020). This wiring algorithm requires developmental flexibility from PNs. The effect is that at the population level, Kenyon cell odor responses change little despite radical alterations to Kenyon cell number. Moreover, we show here that these population-level representations are good enough to allow learned interpretation of molecularly distinct odorants.

Just as PNs adjust to Kenyon cell number, we find that MBONs adjust to Kenyon cell number. MBON activity remained time-locked to odor delivery when Kenyon cell numbers were increased or diminished, and MBONs were able to be activated to a similar level by Kenyon cell repertoires varying by almost an order of magnitude. This suggests that the development of synaptic connectivity between Kenyon cells and MBONs also compensates for changes in Kenyon cell number in a way that maintains coding potential. Previous work on the α’2 MBON showed that this cell can seek input from alternative Kenyon cell partners when its preferred Kenyon cells are lost (Lin et al., 2022). Together, these findings suggest that in developing their dendrites, MBONs seek a particular complement of input and exhibit flexibility in obtaining it. Developmental algorithms that prioritize quantitative relationships across circuit layers prioritize computations and allow variably composed mushroom bodies to detect, discriminate, and drive behavioral responses to molecularly distinct odorants.

In contrast to the developmental accommodation that MBONs make for Kenyon cell number, excess Kenyon cell *activity* in the *Tao* knockdown case propagates forward and produces excess MBON activity in response to some odors. The inability of MBONs to compensate for excess presynaptic activity suggests that sensory input does not regulate Kenyon cell:MBON functional connections. Instead, MBONs could count either Kenyon cell anatomic structures or the Kenyon cell: MBON synapse weight could be scaled by patterned, stimulus-independent neuronal activity (“PSINA”) (Bajar et al., 2022). Sensory drive was also recently shown to be dispensable for PN:Kenyon cell connectivity, as animals lacking the obligate olfactory receptor subunit Orco have PN: Kenyon cell connections that are indiscriminable from controls (Hayashi et al., 2022).

### Effect of *Tao* knockdown on Kenyon cell anatomy and function

Given the developmental robustness with which key computational variables are wired into the mushroom body, altering these variables was a challenge. As we were unable to change Kenyon cell input density by altering Kenyon cell number or the complexity of odor inputs, we sought to directly manipulate Kenyon cell dendritic branching. Different kinds of neurons have diverse dendritic branching patterns, and yet these patterns are specified as an aspect of cell identity and consistent for the same neuron types across animals (Jan and Jan, 2010). We imagine that many, or even all, the gene expression events in a particular cell type will influence dendrite structure, and that the differences between different kinds of neurons could result from myriad transcriptional differences. We have not yet defined the gene regulatory programs that give Kenyon cells their unique claw structures, nor how they count to six. Nevertheless, we found that *Tao* may universally restrain dendritic branching: Just as reducing *Tao* expression in the sensory periphery results in arbor overgrowth, reducing *Tao* in Kenyon cells allows their dendrites to grow more complex (Hu et al., 2020). Indeed, *Tao*-knockdown Kenyon cells seemed to search hard for input, even leaving the calyx in some cases. One area for future research may be to determine if *Tao* levels naturally vary across cells of different dendritic complexity, as do levels of *cut* and *knot* (Jan and Jan, 2010).

### Effect of ectopic *Dscam1* expression on Kenyon cell anatomy and function

In *Drosophila melanogaster*, few proteins have been identified that traffic preferentially to dendrites. One well-studied example, often used to mark dendritic compartments, is the set of Dscam1 isoforms that contain transmembrane domain 1 (Dscam1[TM1]) (Wang et al., 2004). In addition to alternative transmembrane domains, alternative splicing of the Dscam1 ectodomain produces thousands of distinct isoforms; typically, different neurons produce different ectodomains, while common ectodomains on individual neurites of the same neuron induce homophilic repulsion (Hattori et al., 2009; Matthews et al., 2007; Schmucker et al., 2000; Soba et al., 2007). This molecular system allows neurons to avoid synapsing with themselves. In addition to ensuring neuronal self-avoidance, intra-neuronal repulsion mediated by endogenous homophilic Dscam1 interactions likely provides a propulsive force during axon and dendritic development. As Dscam1[TM1] isoforms traffic preferentially to dendrites, while TM2 isoforms traffic to both axons and dendrites, Dscam1[TM1] influences dendritic structure and Dscam1[TM2] influences both axonal and dendritic structure (Wang et al., 2004).

In our experiments, we ectopically expressed a single Dscam1[TM1] cDNA across all Kenyon cells. We assume that when Kenyon cell dendrites all carry the same Dscam ectodomain, inter-neuronal repulsion limits arbor growth (Hattori et al., 2009; Matthews et al., 2007; Soba et al., 2007). In this condition, elongating dendritic neurites of Kenyon cells might experience a terrain similar to that encountered by the train Tootle when he jumps off the tracks: Everywhere he turns, he sees a stop sign (Crampton et al., 1946). While dendritic claw number, number of PN boutons, and calyx size are strongly reduced in this condition, we find that Kenyon cell claws that form appear to make anatomically normal connections with PNs and that Kenyon cells still respond functionally to odors. Because PNs retain endogenous Dscam1 splicing in this condition (i.e. each PN would carry a random set of Dscam1 isoforms, mostly not matching the ectopic isoform we expressed in Kenyon cells), PNs and Kenyon cells would not experience Dscam1-mediated repulsion when contacting one another. Just as the expansion of Kenyon cell dendrites in the *Tao*-knockdown condition prompts PNs to produce more boutons than is typical for that number of Kenyon cells, claw reduction via ectopic Dscam1[TM1] expression results in PNs producing fewer boutons per Kenyon cell. Together, these results suggest that PN bouton development is quantitatively matched to both Kenyon cell number and claw number.

### Claw number determines sparsity of Kenyon cell odor responses

In wild type animals, Kenyon cells are more stringent in their firing than projection neurons, which is thought to allow them to encode glomerular combinations and thus separate olfactory patterns (Gruntman and Turner, 2013; Honegger et al., 2011; Murthy et al., 2008; Turner et al., 2008; Wilson et al., 2004). When we forced Kenyon cells to make fewer claws, a smaller proportion of cells responded to each odor than in wild type, while when we forced Kenyon cells to make more claws, a larger fraction of cells responded to each odor, and some cells responded to all odors. To understand why this would be the case, we considered what active inputs are required for wild type Kenyon cells to spike, and how this threshold behaved when we manipulated claw number. In our prior work, we found that Kenyon cells were very likely to be activated when at least three of their inputs were active, that two active inputs often sufficed, and that one active input could activate a Kenyon cell in certain conditions (Gruntman and Turner, 2013). We considered two models for how Kenyon cell spike threshold would be affected in our manipulations: (A) The number of active claws needed to spike the Kenyon cell is proportional to the total number of claws, or (B) The number of active claws needed to spike the Kenyon cell is constant.

In our previous work, we found that 5-80% of PNs were activated by different odors, with a median of 23% (Gruntman and Turner, 2013). This is consistent with assessment of glomerular responses in the antennal lobe (Knaden et al., 2012). We estimate that for wild type animals, 41% of Kenyon cells with six claws would receive at least two inputs from among the 23% of boutons that are active in response to a median odor, and that 13% of six-clawed Kenyon cells would have at least three active inputs. A spiking threshold of 2-3 claws is therefore consistent with our experimental observation here that 20-40% of Kenyon cells respond to each odor.

*Example calculation for 6-clawed Kenyon cell:*

No inputs active: 0.77^!^ ≈ 0.21

Exactly one input active: 0.23 × 0.77^“^ × _6_C_1_ ≈ 0.37 Exactly two inputs active: 0.23^#^ × 0.77^$^ × _6_C_2_ ≈ 0.28 Exactly three inputs active: 0.23^%^ × 0.77^%^ × _6_C_3_ ≈ 0.11 Exactly four inputs active: 0.23^$^ × 0.77^#^ × _6_C_4_ ≈ 0.02 Exactly five inputs active: 0.23^“^ × 0.77 × _6_C_5_ ≈ 0.003 All six inputs active: 0.23^!^ ≈ 0.0001

At least 2 active ≈0.28 + 0.11 + 0.02 + 0.003 + 0.0001 ≈ 0.41

At least 3 active ≈0.11 + 0.02 + 0.003 + 0.0001 ≈ 0.13

Dscam1[TM1]-overexpressing Kenyon cells had 0-2 claws (median of 1). If the spiking threshold was reduced in these animals, such that only one active claw was required for the Kenyon cell to fire, we would expect ∼23% of Kenyon cells to respond to a median odor, because 23% of boutons would be active. However, we found that odor responses were more dampened than this, with only ∼10% of Kenyon cell responding per odor. This result suggests that the number of active claws needed to spike the Kenyon cell remained the same in this condition, consistent with hypothesis B.

*Tao*-knockdown Kenyon cells had ∼10-14 claws. If the “active claws needed” threshold rises in these animals, such that ∼5 or more active claws were required for the Kenyon cell to fire, we would expect only ∼10% of cells to respond to a median odor (i.e. only 10% of Kenyon cells would have five or more claws innervating the 23% of active boutons). Again, this is inconsistent with what we see. For a Kenyon cell with 12 claws, 53% of cells would have at least three active claws in response to a median odor, and 79% would have at least two active claws. This threshold, the same as for a wild type six-clawed Kenyon cell, is more consistent with our experimental observation that 80% of expanded-claw Kenyon cells in the *Tao*-knockdown condition are activated by each odor. This result is also consistent with hypothesis B—that the number of active claws needed for the Kenyon cell to spike stays constant when claw number is increased due to *Tao* knockdown.

### What have we done to APL?

Kenyon cells activate an inhibitory neuron, APL, that feeds back on Kenyon cells to sparsen their firing (Lin et al., 2014). Recent work suggests that APL provides local feedback inhibition (Amin et al., 2020; Inada et al., 2017; Prisco et al., 2021). Our work here suggests that APL’s activity is determined by the proportion of active Kenyon cells, not the absolute number of active Kenyon cells. If APL was responsive to the absolute number of active Kenyon cells, it would provide excess inhibition in brains with more Kenyon cells and limited inhibition in brains with few Kenyon cells; this relationship would produce the same effect as our rejected model A above, where Kenyon cells become less responsive to odors as claw number grows. We do not see reduction in Kenyon cell activity or MBON responses when Kenyon cell number or claw number rises, nor increases in Kenyon cell activity or MBON responses when Kenyon cell number falls. We therefore predict that APL makes the same number of synapses in hemispheres with 500, 2000, or 4000 Kenyon cells, such that it gets less synapses per Kenyon cell in a brain with more Kenyon cells and vice versa. This would allow APL to provide feedback inhibition that is proportional to Kenyon cell number. Alternatively or in addition to such a developmental scaling mechanism, homeostatic mechanisms could adjust Kenyon cell:APL coupling in adulthood such that Kenyon cell repertoires of different sizes evoke comparable APL responses (Apostolopoulou and Lin, 2020).

### Quantitative variables of the mushroom body expansion layer are developmentally independent

Expansion layers, in which a particular number of sensory input channels is mapped onto a much larger set of second-order neurons, are thought to increase the number of stimuli that an animal can detect and discriminate, i.e. the dimensionality of the representation. Many wiring and functional parameters influence the dimension of the representation that second-order neurons can extract from a fixed number of sensory channels. These parameters include the number of second-order neurons, the number of inputs each second-order neuron receives, and the “expansion ratio,” which is the relationship between these two variables. The number of inputs that need to be active for the second-order neuron to fire and the strength of feedback inhibition are also expected to strongly influence odor coding. In theoretical work, these parameters can be dissociated and freely varied. We will discuss how these variables behave developmentally using the framework provided by (Litwin-Kumar et al., 2017). A summary of these variables and how we have altered them here is provided in Supplemental Tables 1 and 2.

Our results point to a distinction between what is mathematically possible and what is biologically plausible. In particular, two key parameters limiting the dimension of Kenyon cell odor responses behave differently in our developmental manipulations than how they are typically manipulated in a model: (1) Claw number and Kenyon cell number are not linked in our results, (2) nor are claw number and claw strength. For maximizing dimensionality of odor responses, more expansion layer neurons would almost always be better, so defining the ideal number of inputs per cell in a model depends on creating a constraint where the total number of PN-Kenyon cell connections (approximated by Kenyon cell number * claw number) stays constant. We initially expected to find a tradeoff like this developmentally: i.e. we expected that if we increased Kenyon cell number, the claw number of individual cells would go down and vice versa. However, we found that because Kenyon cells keep their claw numbers constant and developmentally instruct PNs, the total connections in the calyx rises and falls with the number of Kenyon cells. Similarly, when we forced increases and decreases in claw number per Kenyon cell, we saw only small changes in Kenyon cell number that could not compensate for claw increase. Therefore, total connections in the calyx changed as we changed either the number of Kenyon cells or the number of inputs per Kenyon cell.

We can imagine cell number and claw number as being independently tuned during evolution: selection may operate on the one hand to balance the total size and influence of the mushroom body with respect to the rest of the brain--consistent with the importance of learned versus innate decision-making to the survival of different types of animals--and on the other hand to determine input number per cell. Second, selection may operate not to maximize perceptual discrimination (which will always be better with more expansion layer neurons, as indeed we observed experimentally here) but to balance dimensionality with the need to generalize. For example, a recent comparative analysis suggests that expansion layer neuron number is matched to the complexity of the incoming sensory representation, and hypothesizes that increasing expansion ratio further results in over-fitting (Hiratani and Latham, 2022). Thus while more neurons would always improve dimensionality, it does not necessarily improve decision-making.

Indeed, both the complexity of sensory inputs to the mushroom body and the size of the Kenyon cell population vary by orders of magnitude across arthropods: While flies have ∼2000 Kenyon cells per hemisphere, bees and cockroaches have ∼200,000; the number of olfactory channels varies from 20 to 400; and the mushroom body can receive input from diverse sensory modalities (Brand et al., 2018; Fahrbach, 2006; Farris and Sinakevitch, 2003; Missbach et al., 2014; Witthöft, 1967; Zhao et al., 2008). For example, chemosensation is thought to be an evolutionarily ancient modality, and the mushroom body to have evolved as a chemosensory structure; however, many arthropods have evolved visual learning, and devote as much as half the mushroom body to vision (Farris and Sinakevitch, 2003; Gronenberg, 2001; Wolff and Strausfeld, 2015). In many insects, Kenyon cells even continue to be generated in adulthood (reviewed in (Simões and Rhiner, 2017)).

Just as Kenyon cell number and claw number could be independently varied in our experiments, we infer that directly changing claw number did not influence claw strength nor the number of active inputs needed for the Kenyon cell to spike. The logical manipulation in a model would be to test how distributing the same number of input synapses across different numbers of claws would affect the dimensionality of odor responses, or to vary Kenyon cell spike threshold with input number. However, it appears that these variables are uncoupled developmentally: Biologically manipulating claw number had the effect of changing the total input each Kenyon cell received while leaving the spike threshold the same, such that cells with more claws simply become more promiscuous. These results highlight that the evolutionary optimization of combinatorial coding requires a precise relationship between the dendrite structure of expansion layer neurons and their spike threshold.

Thus, because an increase or decrease in the total number of Kenyon cell claws in the calyx can instruct a change in the total presynaptic sites produced by PNs, claw number and weight do no influence each other in these manipulations, and Kenyon cells with more claws become more promiscuous. However, we expect that individual Kenyon cells competing for input from the available PN presynapse repertoire may indeed balance claw number and strength. For example, within a single brain, individual Kenyon cells with more claws have been shown to have fewer synapses per claw and may require more active inputs to spike (Eichler et al., 2017; Gruntman and Turner, 2013). There is at this time little research examining how and whether neurons “know” what amount of input they should obtain during development, but the rich repertoire of known homeostatic mechanisms that adjust input-output functions in development and adulthood suggest such cellular expectations exist (Goel et al., 2019; Parrish et al., 2014; Tatavarty et al., 2020).

### Effects of manipulating expansion layer neurons on mushroom body outputs and behavior

By testing the two-choice olfactory learning behavior of individual animals, we were able to relate continuous variation in Kenyon cell features to performance for the first time. We found that animals were able to make learned associations and discriminate molecularly distinct odors with only 500 Kenyon cells, versus 4000 cells across hemispheres in controls. In animals with enhanced Kenyon cell repertoires, we found that learning performance improved in larger calyces. In both cases, mushroom body output neuron activity in response to odors roughly matches that of control animals. These results suggest that while animals *can* discriminate molecularly distinct odorants with only a fraction of the normal set of Kenyon cells, the fidelity of their resulting decisions can be improved by increasing the size of the Kenyon cell population. Determining what aspect of discrimination or decision-making underlies this improvement would require monitoring synaptic weight changes in the Kenyon cell output synapse upon olfactory conditioning across animals with variably constituted mushroom bodies.

When we knocked down *Tao* in Kenyon cells, claw number increased by 50%, projection of Kenyon cells to the *α*β and *α*’β’ lobes was disrupted, and Kenyon cell odor responses broadened, such that most cells responded to multiple odors. Moreover, the excess Kenyon cell activity propagated to the γ-lobe MBON in our functional imaging. Even in this extreme case, animals performed two-choice odor learning like champs. Together, our results impress upon us the remarkable functional and developmental robustness of the mushroom body circuit, which continues to compute useful information about the world across a range of hard-wired architectures. Developmental algorithms that prioritize functional computations may allow circuit formation to correct for both acute mistakes or environmental fluctuations and facilitate evolutionary change to wiring parameters.

## Methods

KEY RESOURCES TABLE

KEY RESOURCES TABLE
| REAGENT or RESOURCE | SOURCE | IDENTIFIER |
| --- | --- | --- |
| **Antibodies** |  |  |
| anti-ChAT (mouse monoclonal) | Developmental Studies Hybridoma Bank (DSHB) | 9E10 |
| anti-DsRed (rabbit polyclonal) | Clontech | 632496 |
| anti-GFP (sheep polyclonal) | Bio-Rad | 4745-1051 |
| anti-GFP (chicken polyclonal) Umich (also used to stain mAmetrine, GCaMP, C3PA) | Cai Lab |  |
| anti-GFP (chicken) Janelia | Fisher | A-10262 |
| CyTM3 anti-rabbit (goat) | Jackson Immunoresearch | 111-165-144 |
| CyTM5-anti-mouse (goat) | Jackson Immunoresearch | 115-175-166 |
| Alexa 488 anti-chicken (goat polyclonal) Janelia | Fisher | A32931 |
| Alexa 488 anti-mouse (goat polyclonal) | Fisher | A11029 |
| Alexa 647 anti-mouse (goat polyclonal) | Fisher | A21236 |
| Alexa 488 anti-sheep (donkey polyclonal) | Fisher | A11015 |
| Alexa 488 anti-chicken (goat polyclonal) Umich | Fisher | A11039 |
| Alexa 568 anti-rabbit (goat polyclonal) | Fisher | A11036 |
| anti-brp (mouse monoclonal) | DSHB | Nc82 |
| anti-mTFP (rat) | Cai Lab |  |
| anti-mNeonGreen (rabbit) | Cai Lab |  |
| anti-mKusabiraOrange2 (mouse) | Cai Lab |  |
| anti-mKate2 (guinea pig) | Cai Lab |  |
| Alexa 594 anti-rat (donkey polyclonal) | Fisher | A21209 |
| CF 555 anti-mouse (donkey polyclonal) | Sigma | SAB4600060 |
| CF 633 ant-guinea pig (donkey polyclonal) | Sigma | SAB4600129 |
| Atto490 LS anti-rabbit | Hypermol | 2309 |
| **Chemicals, peptides, and recombinant proteins** |  |  |
| Paraformaldehyde (Clowney Lab) | EMS | 15710 |
| Paraformaldehyde (Janelia) | EMS | 15713 |
| Paraformaldehyde (Cai Lab) | Sigma | P6148 |
| Hydroxyurea | Sigma | H8627 |
| Mineral oil | Sigma | 330760 |
| Paraffin oil | Sigma | 18512 |
| Benzaldehyde | Sigma | 418099 |
| Isobutyl acetate | Sigma | 537470 |
| Ethyl acetate | Sigma | 270989 |
| Methylcyclohexanol | Sigma | 153095 |
| 3-Octanol | Sigma | 218405 |
| Phalloidin-568 (used in stainings, though not shown in figures) | Fisher | A12380 |
| DAPI (used in stainings, though not shown in figures) | Sigma | D9542 |
| Retinal | Janelia Media Facility |  |
| 10x PBS Gibco | Fisher | 70-011-044 |
| Glycerol | Fisher | G33-500 |
| Propyl gallate | Sigma | 02370 |
| Vectashield | Vector Laboratories | H-1000 |
| StartingBlock | Thermo | 37578 |
| Normal goat serum (Umich) | MP biobiomedicals | 8642921 |
| Normal goat serum (Umich) | Abcam | ab7481 |
| Triton | Sigma | X100 |
| Sodium chloride NaCl | Sigma | S7653 |
| Potassium chloride KCl | Fisher | P217-500 |
| Calcium chloride CaCl<sub>2</sub>*2H<sub>2</sub>O | Fisher | C70-500 |
| Magnesium chloride MgCl<sub>2</sub>*6H<sub>2</sub>O | Sigma | M2670 |
| Sodium bicarbonate NaHCO<sub>3</sub> | Fisher | S233-500 |
| Sodium phosphate NaH<sub>2</sub>PO<sub>4</sub>*H<sub>2</sub>O | Fisher | S369-500 |
| Trehalose | Sigma | T9531 |
| Sucrose | Sigma | 84097 |
| HEPES sodium salt | Sigma | H7006 |
| Schneider's insect medium (Janelia) | Sigma | S01416 |
| PBS (Janelia) | Cellgro | 21-040 |
| Normal goat serum (Janelia) | Fisher | 16210-064 |
| Poly-L-lysine hydrobromide | Sigma | P1524 |
| Kodak Photo-Flo 200 Solution (for poly-L-lysine-coating cover slips) | EMS | 74257 |
| DPX Mountant for Microscopy | EMS | 13512 |
| Ethanol. ACS reagent, >99.5% (200 proof) | Sigma | 459844 |
| Xylenes | Fisher | X5-500 |
| **Experimental models: Organisms/strains** |  |  |
| UAS-CD8GFP (II) | BDSC 5130 | (Lee and Luo, 1999) |
| GH146<sup>QF</sup>, QUAS-mtdTomato-3xHA (II) | BL30037 | (Potter et al., 2010) |
| P(caryP) (attp2) | BL36303 | (Perkins et al., 2015) |
| OK107<sup>Gal4</sup> (IV) | BL 854 | (Connolly et al., 1996) |
| UAS-mud-RNAi (attp2) | BL35044 | (Perkins et al., 2015) |
| UAS-GCaMP6s (attp40) | BL 42746 | (Akerboom et al., 2012) |
| UAS-GCaMP6s (VK0005) | BL 42749 | (Akerboom et al., 2012) |
| UAS-Tao1-RNAi (attp2) | BL 35147 | (Perkins et al., 2015) |
| GMR58E02-lexA (attP40) | BL 52740 | Pfeiffer, Rubin<br>LexA Collection |
| MZ19<sup>QF</sup> (II) | BL41573 | (Hong et al., 2012) |
| QUAS-mtdTomato-3xHA (II) | BL 30004 | (Potter et al., 2010) |
| 58F02-Gal4 (attP2) | BL 39186 | (Jenett et al., 2012) |
| UAS-CD8GFP (III) | BL 5130 | (Lee and Luo, 1999) |
| 10XUAS-IVS-myr::GFP (attP2) | BL 32197 | (Pfeiffer et al., 2010) |
| R25D01-lexA (attP40) | BL 53519 | Pfeiffer, Rubin<br>LexA Collection |
| 13XLexAop2-IVS-GCaMP6s-SV40 (su(Hw)attP5) | BL 44589 | FBrf0221946 |
| 10XUAS-IVS-myr::tdTomato (attP2) | BL 32221 | (Pfeiffer et al., 2010) |
| R53C03-lexA (attP40) | BL 52861 | Pfeiffer, Rubin<br>LexA Collection |
| UAS-Dscam<sup>3.36.25.1</sup>::RFP on II | via Vanessa Ruta | Tzumin Lee lab,<br>made by Jacob<br>Yang following<br>(Yu et al., 2009) |
| LexAopChrimson::tdTomato on III | BL 55139 |  |
| Gr64f-LexA on III |  | (Rajagopalan et al., 2022) |
| P{hs-KD}attP3 | BL 56167 | (Awasaki et al., 2014) |
| mBitbow 2.2 (mem5FP KDon>FLP>) VK00027 |  | This paper |
| UAS-Dscam<sup>3.36.25.1</sup>::GFP on X | via Bing Ye | (Wang et al., 2004) |
| LexAopChrimson::tdTomato on X | Unknown | Janelia fly facility |
| **Software and algorithms** |  |  |
| Suite2p |  | (Pachitariu et al., 2017) |
| FIJI |  | Schindelin et al., 2012 |
| FIJI plugin “Blind Analysis Tools” |  | https://imagej.net/Blind_Analysis_Tools |
| Cellpose |  | Stringer et al., 2021 |
| GraphPad Prism 9 |  |  |
| R |  |  |
| R package “corrplot” |  |  |
| Python |  |  |
| MATLAB 2018b, Mathworks |  |  |
| MATLAB 2020b, Mathworks |  |  |
| **Other** |  |  |
| Duck EZ Start Packaging tape | Office Depot | 511879 |
| UV Glue | Loctite | 3106 |
| JAXMAN 365nm Flashlight | Amazon | B077GPXBK1 |
| Electra Waxer | Almore | 66000 |
| Gulfwax Paraffin | Grocery store | C0130 |
| Hair | EJC |  |
| Umich fly food | Lab Express, Ann Arbor, MI | “B” food |
| Janelia fly food |  |  |
| Heat packs for mailing flies | TSKSupply | 40hour Uniheat |
| Thermometers for mailing flies | ThermoPro | TP50 |
| Cell strainer caps (used as colanders for stainings) | Fisher | BD 352235 |
| Binder reinforcement stickers (for mounting brains) | Office Depot | Avery 5722 |
| Terasaki plates Greiner Bio-One (for staining) | Fisher | 07-000-623 |
| Grape juice agar plates | Lab Express, Ann Arbor, MI |  |
| Coffee filters | Departmental coffee supplies |  |
| Caps for odor vials | Sigma | 27020 |
| Odor vials | Sigma | 27088-U |
| Cover glass 22x22 mm Square No. 1. Corning. # (for mounting, Janelia) | Corning | CLS284522-2000EA |
| Cover glass 22x22 mm Square No. 2. (for spacers, Janelia) | Fisher | # 12-540B |
| Cover glass staining jar (Janelia) | EMS | 72242-24 |
| Diamond Cleaver (Janelia) | Diatome.com | CLE |
| DPX Slide Jig | Janelia Instrument Design & Fabrication | http://hhmi.flintbox.com/public/project/26605 |
| Drosophila Mounting T-dish. 8x10 (Janelia, for adult brains) | Janelia Instrument Design & Fabrication | http://hhmi.flintbox.com/public/project/26606 |
| Superfrost Plus Slides 25x75 mm | Fisher | 12-550-15 |

## EXPERIMENTAL MODEL AND SUBJECT DETAILS

### Flies

Flies used for anatomy and functional imaging were maintained on cornmeal (‘Bloomington-B’) food (Lab Express, Ann Arbor, MI) augmented with a yeast sprinkle in a humidified incubator at 25°C on a 12:12 light:dark cycle. Flies for dense Bitbow labeling (Figure 2A) were heat shocked at the 1^st^ instar larval stage for 30 mins in a 37°C water bath, and the adults were collected 4 days after eclosion for immunostaining and imaging.

Flies used for behavioral experiments were maintained on Janelia Research Campus’s standard cornmeal food supplemented with 0.2 mM all-trans-retinal. HU ablation flies and sham controls were reared in a humidified incubator at 21°C on a 12:12 light:dark cycle, while all other flies were reared at 25°C on a 12:12 light:dark cycle. Flies were kept in the dark throughout.

Experimental crosses for ablation experiments were set up with large numbers of males (60-100 per cage) and females (300-500 per cage). All other behavior crosses were set up with 8 male and 8 female parents and a sprinkle of yeast was added to the food vials. Cross progeny (2-7 days old) were sorted on a cold plate at around 4°C and females of the appropriate genotype were transferred to starvation vials. Starvation vials contained nutrient-free 1% agarose to prevent desiccation. Flies were starved between 36-60 hrs before being aspirated into the Y-arena for experiments.

### Genotypes used in each figure

|  |  |
| --- | --- |
| Figure 1A | UAS-CD8GFP/+; GH146<sup>QF</sup>, QUAS-mtdTomato-3xHA/P(caryP)attp2; OK107<sup>Gal4</sup>/+<br>male, 2-3 days old |
| Figure 1C | LexAopChrimson::tdTomato/+; Gr64f-LexA/UAS-mud RNAi; OK107<sup>Gal4</sup>/+<br>female, 3-7 days old |
|  | LexAopChrimson::tdTomato/+; Gr64f-LexA/P(caryP)attp2; OK107<sup>Gal4</sup>/+<br>female, 3-7 days old |
| Figure 1E-J, S1 | UAS-GCaMP6s/+; UAS-GCaMP6s/UAS-mud RNAi; OK107<sup>Gal4</sup>/+<br>mix of males and females, 2-5 days old |
|  | UAS-GCaMP6s/+; UAS-mud RNAi/TM2; OK107<sup>Gal4</sup>/+<br>mix of males and females, 2-5 days old |
|  | UAS-GCaMP6s/+; UAS-GCaMP6s/P(caryP)attp2; OK107<sup>Gal4</sup>/+<br>mix of males and females, 2-5 days old |
|  | UAS-GCaMP6s/+; P(caryP)attp2/TM2; OK107<sup>Gal4</sup>/+<br>mix of males and females, 2-5 days old |
| Figure 2C, D, S2B | hsKD; CyO/+; Bitbow 2.2/UAS-Tao RNAi; OK107<sup>Gal4</sup>/+<br>males, 2-5 days old |
|  | hsKD; CyO/+; Bitbow 2.2/+; OK107<sup>Gal4</sup>/+<br>males, 2-5 days old |
| Figure 2E-G, 2I-N | UAS-CD8GFP/+; GH146<sup>QF</sup>, QUAS-mtdTomato-3xHA/UAS-Tao RNAi; OK107<sup>Gal4</sup>/+<br>males, 3 days old |
|  | UAS-CD8GFP/+; GH146<sup>QF</sup>, QUAS-mtdTomato-3xHA/P(caryP)attp2; OK107<sup>Gal4</sup>/+<br>males, 3 days old |
| Figure 2H | UAS-mCD8GFP/+; Gr64f-LexA, LexAop cs-Chrimson-tdTomato/P(caryP)attp2; OK107<sup>Gal4</sup>/+<br>females, 3-7 days old |
|  | UAS-mCD8GFP/+; Gr64f-LexA, LexAopcs-Chrimson-tdTomato/UAS-Tao RNAi; OK107<sup>Gal4</sup>/+<br>females, 3-7 days old |
| Figure 3, S3 | UAS-GCaMP6s/+; UAS-GCaMP6s/UAS-Tao RNAi; OK107<sup>Gal4</sup>/+<br>mix of males and females, 2-5 days old |
|  | UAS-GCaMP6s/+; UAS-Tao RNAi/TM2; OK107<sup>Gal4</sup>/+<br>mix of males and females, 2-5 days old |
|  | UAS-GCaMP6s/+; UAS-GCaMP6s/P(caryP)attp2; OK107<sup>Gal4</sup>/+<br>mix of males and females, 2-5 days old |
|  | UAS-GCaMP6s/+; P(caryP)attp2/TM2; OK107<sup>Gal4</sup>/+<br>mix of males and females, 2-5 days old |
| Figure 4A-B | hsKD; Dscam<sup>3.36.25.1</sup>:RFP/Bl; Bitbow 2.2/+; OK107<sup>Gal4</sup>/+<br>males, 3-4 days old |
|  | hsKD; Bl/+; Bitbow 2.2/+; OK107<sup>Gal4</sup>/+<br>males, 3-4 days old |
| Figure 4C-K | Dscam<sup>3.36.25.1</sup>:GFP; UAS-CD8GFP/Pin; GH146<sup>QF</sup>, QUAS-mtdTomato-3xHA/+; OK107<sup>Gal4</sup>/+<br>males, 2-3 days old |
|  | Dscam<sup>3.36.25.1</sup>:GFP; Pin/CyO; TM2/+; OK107<sup>Gal4</sup>/+<br>males, 2-3 days old |
|  | UAS-CD8GFP/Sp; GH146<sup>QF</sup>, QUAS-mtdTomato-3xHA/TM2; OK107<sup>Gal4</sup>/+<br>males, 2-3 days old |
|  | Bl/CyO; TM2/TM6B; OK107<sup>Gal4</sup>/+<br>males, 2-3 days old |
| Figure 5, S5 | UAS-GCaMP6s/Dscam<sup>3.36.25.1</sup>:RFP; UAS-GCaMP6s/+; OK107<sup>Gal4</sup>/+<br>mix of males and females, 2-5 days old |
|  | UAS-GCaMP6s/Dscam<sup>3.36.25.1</sup>:RFP; UAS-GCaMP6s/TM2; OK107<sup>Gal4</sup>/+<br>mix of males and females, 2-5 days old |
|  | Males: UAS-GCaMP6s/+; UAS-GCaMP6s/+; OK107<sup>Gal4</sup>/+<br>Females: yw/+; UAS-GCaMP6s/+; UAS-GCaMP6s/+; OK107<sup>Gal4</sup>/+<br>2-5 days old |
|  | Males: UAS-GCaMP6s/+; UAS-GCaMP6s/TM2; OK107<sup>Gal4</sup>/+<br>Females: yw/+; UAS-GCaMP6s/+; UAS-GCaMP6s/TM2; OK107<sup>Gal4</sup>/+<br>2-5 days old |
| Figure 6B-F, S6A-E | 13x LexAop cs-Chrimson-tdTomato/+; 58E02-LexA/Mz19<sup>QF</sup>, QUAS-mtdTomato-3xHA; 58F02-Gal4, UAS-mCD8GFP/10xUAS-IVS-myrGFP<br>females, 3-8 days old |
| Figure 6G-I, S6F,G | R25D01-LexA, LexAop-GCaMP6s/CyO; UAS-tdTomato/TM6B; OK107<sup>Gal4</sup>/OK107<sup>Gal4</sup><br>mix of males and females, 2-5 days old |
| Figure 7A-D | UAS-mCD8GFP/+; Gr64f-LexA, LexAop cs-Chrimson-tdTomato/P(caryP)attp2; OK107<sup>Gal4</sup>/+<br>females, 3-7 days old |
|  | UAS-mCD8GFP/+; Gr64f-LexA, LexAop cs-Chrimson-tdTomato/UAS-mud RNAi; OK107<sup>Gal4</sup>/+<br>females, 3-7 days old |
|  | UAS-mCD8GFP/+; Gr64f-LexA, LexAop cs-Chrimson-tdTomato/UAS-Tao RNAi; OK107<sup>Gal4</sup>/+<br>females, 3-7 days old |
| Figure 7E-G, S7A,B | R25D01-LexA, LexAop-GCaMP6s/+; UAS-tdTomato/P(caryP)attp2; OK107<sup>Gal4</sup>/+<br>mix of males and females, 2-5 days old |
|  | R25D01-LexA, LexAop-GCaMP6s/+; UAS-tdTomato/UAS-mud RNAi; OK107<sup>Gal4</sup>/+<br>mix of males and females, 2-5 days old |
|  | R25D01-LexA, LexAop-GCaMP6s/+; UAS-tdTomato/UAS-Tao RNAi; OK107<sup>Gal4</sup>/+<br>mix of males and females, 2-5 days old |

While we used mixed-sex cohorts when possible, certain genetic and experimental approaches required us to use one sex or the other, as detailed below under behavioral and image analysis (e.g. males are too small for the Y-arena).

## METHOD DETAILS

### Immunostainings

This protocol is adapted from (Elkahlah et al., 2020), with some modifications. Brains were dissected for up to 20 min in AHL dissection saline (108 mM NaCl, 5 mM KCl, 2 mM CaCl2, 8.2 mM MgCl2, 4 mM NaHCO3, 1 mM NaH2PO4, 5 mM trehalose, 10 mM sucrose, 5 mM HEPES pH7.5, osmolarity adjusted to 265 mOsm). For most experiments, brains were transferred to 1% paraformaldehyde in PBS on ice. All steps were performed in cell strainer baskets (caps of FACS tubes) in 24 well plates, with the brains in the baskets lifted from well to well to change solutions. Brains were fixed overnight at 4C in 1% PFA in PBS. On day 2, brains were washed 3 × 10’ in PBS supplemented with 0.1% triton-x-100 on a shaker at room temperature, blocked 1 hr in PBS, 0.1% triton, 4% Normal Goat Serum, and then incubated for at least two overnights in primary antibody solution, diluted in PBS, 0.1% triton, 4% Normal Goat Serum. Primary antibody was washed 3 × 10’ in PBS supplemented with 0.1% triton-x-100 on a shaker at room temperature, then brains were incubated in secondary antibodies for at least two overnights, diluted in PBS, 0.1% triton, 4% Normal Goat Serum. When used, DAPI (one microgram/mL) and phalloidin 568 (1:40-1:80) were included in secondary antibody mixes. Primary antibodies used were mouse anti-ChAT 9E10 (1:200, DSHB), rabbit anti-dsRed (1:500-1:1000, Clontech), sheep anti-GFP (1:250-1:1000, Bio-Rad), chicken anti-GFP (1:5000, Dawen Cai), mouse anti-nc82 (1:30, DSHB). Secondary antibodies were Alexa 488, 568, and 647 conjugates (1:500, Invitrogen). Brains were mounted in 1x PBS, 90% glycerol supplemented with propyl gallate in binder reinforcement stickers sandwiched between two coverslips. Samples were stored at 4°C in the dark prior to imaging. The coverslip sandwiches were taped to slides, allowing us to perform confocal imaging on one side of the brain and then flip over the sandwich to allow a clear view of the other side of the brain. This allowed us to score features on the anterior and posterior sides of each sample. Scanning confocal stacks were collected along the anterior-posterior axis on a Leica SP8 with one micrometer spacing in Z and ∼200 nm axial resolution.

For sparse Bitbow labeling (Figures 2C-D, 5I-J), brains were fixed in 4% paraformaldehyde at room temperature for 30 minutes. This was followed by washing 3 × 10’ in PBS supplemented with 0.1% triton-x-100 on a shaker at room temperature. The brains were blocked 1 hr in PBS, 0.1% triton, 4% Normal Goat Serum or StartingBlock. All the steps were carried out in Terasaki plates, with each well containing 2-3 brains to keep track of fixation, wash, and block timing.

The brains were then moved to 24-well plates for incubation with primary antibody solution at 4C. Subsequent steps of secondary antibody incubation, mounting and imaging were the same as described above. Primary antibodies were rat anti-mTFP (1:500), chicken anti-GFP (1:5000), rabbit anti-mNeonGreen (1:500), mouse anti-mKusabira-Orange2 (1:500), guinea pig anti-mKate2 (1:500) (Dawen Cai Lab) and secondary antibodies were Alexa 488, 594, 647 conjugates (1:500, Invitrogen), CF 555, CF 633 (1:500, Sigma), and Atto490 LS (1:250, Hypermol). For dense Bitbow labeling in Figure 2A, a similar protocol was followed as described in Li et al., 2021. The difference from the sparse Bitbow protocol above was that the mounting media was Vectashield. The confocal image was acquired with Zeiss LSM780 with a 40 × 1.3 NA oil immersion objective (421762-9900-000). A 32-channel GaAsP array detector was used to allow multi-track detection of five fluorophores with proper channel collection setups.

The following protocol applies to the dissections, immunohistochemistry and imaging of fly brains done post-behavior (Figures 6, 7), and for the Brp staining (Figure 1D, 2H). These were done principally as previously described ((Aso et al., 2014a); https://www.janelia.org/project-team/flylight/protocols). In brief, brains were dissected in Schneider’s insect medium and fixed in 2% paraformaldehyde (diluted in the same medium) at room temperature for 55 mins.

Tissues were washed in PBT (0.5% Triton X-100 in phosphate buffered saline) and blocked using 5% normal goat serum before incubation with antibodies. Since flies had to be tracked individually, single dissected tissues were fixed and washed in microwell plates. After washes in the microwell plate, tissues were mounted on poly-L-lysine-coated cover slips before performing immunohistochemistry, post-fixation, and embedding in DPX as described. Tissues expressing GFP and tdTomato were incubated with chicken anti-GFP (1:1000, Fisher) together with rabbit anti-dsRed (1:1000, Clontech), and mouse anti-BRP hybridoma supernatant (1:30, DSHB), followed by Alexa Fluor 488-conjugated goat anti-chicken (Fisher), CyTM3-conjugated goat anti-rabbit (Jackson ImmunoResearch), and CyTM5-conjugated goat anti-mouse (Jackson ImmunoResearch). Some tissues were labeled with mouse anti-ChAT hybridoma supernatant (1:100, DSHB) instead of anti-BRP, while all other antibodies remained the same. Nuclei were stained with DAPI. Image z-stacks were collected using an LSM880 confocal microscope (Zeiss, Germany) fitted with Plan-Apochromat 40X/1.3 and Plan-Apochromat 63X/1.40 oil immersion objectives. All tissues were first imaged at 40X with a voxel size of 0.44 x 0.44 x 0.44 mm3 (1024 x 1024 pixels in xy). Selected tissues were imaged at higher resolution in one of three ways: at 40X with a voxel size of 0.22 x 0.22 x 0.44 mm3 (2048 x 2048 pixels in xy; “40X hi-res”), at 63X with a voxel size of 0.19 x 0.19 x 0.37 mm3 (1024 x 1024 pixels in xy; “63X hi-res-1”), or at 63X with a voxel size of 0.1 x 0.1 x 0.37 mm3 (2048 x 2048 pixels in xy; “63X hi-res-2”).

### Molecular cloning of mBitbow2.2

DNA fragment of EcoRI-FrtF15-KDRT-FlpN-KDRT-SwaI was synthesized by GeneArt (Thermo Fisher), and cloned in pattB-nSyb-sr(Flp)F15 (Li et al., 2021) to replace the N-terinal region of flippase, yielding pattB-nSyb-sr(KDonFlp)F15. nSyb-sr(KDonFlp)F15 was then amplified by PCR and integrated into NdeI-linearized mBitbow1.0 through Gibson assembly, yielding mBitbow2.2.

### HU ablation

To generate flies with reduced Kenyon cell numbers, we used an existing chemical method to ablate KC neuroblasts (de Belle and Heisenberg, 1994; Elkahlah et al., 2020; Sweeney et al., 2012). In *Drosophila*, 4 mushroom body neuroblasts from each hemisphere give rise to ∼500 KCs each (Ito et al., 1997). Most neuroblasts pause their divisions during the first 8 hours after larval hatching (ALH), however MB neuroblasts continue to divide. Therefore, by feeding larvae a mitotic poison hydroxyurea (HU) during this time window, Kenyon cell neuroblasts can specifically be ablated (de Belle and Heisenberg, 1994). We can achieve a wide range of KC numbers by tweaking the concentration and duration of HU application (Elkahlah et al., 2020). This allowed us to generate flies with 0 to 4 KC neuroblasts, and the effect on each hemisphere was independent of the other resulting in a large range of KC numbers. However, as we described previously, the most informative brains, with intermediate numbers of KC neuroblasts remaining, are difficult to obtain—most ablation batches have a preponderance of unaffected or fully ablated mushroom bodies.

The protocol was adapted from (Elkahlah et al., 2020; Sweeney et al., 2012). We set up large populations of flies in cages two days prior to ablation and placed a 35 or 60 mm grape juice agar plate (Lab Express, Ann Arbor, MI) in the cage with a dollop of goopy yeast. One day prior to the ablation, we replaced the grape juice/yeast plate with a new grape juice/yeast plate. On the morning of the ablation, we removed the plate from the cage and discarded the yeast puck and any hatched larvae on the agar. We then monitored the plate for up to four hours, until many larvae had hatched. Larvae were washed off the plate using a sucrose solution, and eggs were discarded. Larvae were then strained in coffee filters, and submerged in hydroxyurea (Sigma, H8627) in a yeast:AHL mixture, or sham mixture without HU. Ablation condition was 10 mg/mL HU, given for 1 hour. One batch experienced 15 mg/mL HU, given for 1 hour; this was done to gather more data points with a lower KC clone count. Larvae were then strained through coffee filters again, rinsed, and placed in a vial or bottle of B food (for MBON functional imaging and immunohistochemistry) or Janelia food supplemented with 0.2 mM all-trans-retinal (for behavior) until eclosion. We opened a new container of hydroxyurea each month as it degrades in contact with moisture, and we found its potency gradually declined.

We achieved a U-shaped distribution of the HU effect, with many samples unaffected and many with all four KC neuroblasts lost. Ablated animals along with the control group (sham) were shipped in temperature-controlled conditions to Janelia Research Campus for behavior. A digital thermometer was kept in each shipment to record the lowest and highest temperature experienced during shipping. Batches that experienced ∼15-27 Celsius were used for experiments. Animals were either shipped as larvae or late pupae, and not as adults in order to give enough time window for using the appropriate age of adult flies for behavior experiments.

### Identification of driver lines for relevant cell populations

LexA drivers for MBONs were screened from Janelia FlyLight online collection database and previous split-GAL4 characterization (Aso et al., 2014a). Among the LexA drivers, R25D01 was selected based on its specific expression in γ2,α’1 MBON, involvement in odor-associated appetitive memory formation, and ability to drive quantifiable GCaMP6s expression (Aso et al., 2014b; Handler et al., 2019). We used R25D01-LexA to drive expression of GCaMP6s in γ2,α’1 MBON for MBON functional imaging, and to assess connection between KCs and MBONs morphologically.

### *In vivo* functional imaging

Protocol was adapted from (Ruta et al., 2010) and (Elkahlah et al., 2020). We prepared 2–5 day old adult flies for *in vivo* two photon calcium imaging on a Bruker Investigator, affixing the fly to packaging tape (Duck EZ Start) with human hair and UV glue (Loctite 3106). The fly was tilted to allow optical access to KC somata, at the dorsal posterior surface of the brain. For MBON axonal imaging, the fly was similarly prepped to allow optical access to the mushroom body lobes for visualizing MBON axons; these flies also carried a Kenyon cell label to serve as an anatomical guide. We waxed the proboscis in an extended position to reduce motion.

Imaging was performed in external saline (AHL). For odor delivery, half of a 1200 mL/min airstream, or 600 mL/min, was directed toward the antennae through a Pasteur pipette mounted on a micromanipulator. At a trigger, 10% of the total air stream was re-directed from a 10 mL glass vial containing mineral oil to a vial containing odorants diluted 1:10 in mineral oil, or a second vial of mineral oil (mechanosensory control). Final odor dilution was therefore 1:100. Filter paper ‘wicks’ were inserted into each vial to allow odor to saturate the vial headspace. Odors were delivered for two seconds, with 30 s in between stimulations. We used a simplified olfactometer capable of delivering five different odorants in which overall airflow was metered by analogue flowmeters (Brooks Instruments) and valve switching controlled by an Arduino. Odor delivery was initially optimized using a mini-PID placed in the half of the air stream not directed at the fly (Aurora Biosciences). Images were collected at ∼5 Hertz, and we imaged a single plane for each sample. Odors used were ethyl acetate (EA), isobutyl acetate (IBA), benzaldehyde (BZH), octanol (OCT), and methylcyclohexanol (MCH). Recording neuronal responses to the mineral oil control allows us to monitor sensory components that are a shared across the different odor stimuli, including olfactory responses to volatiles in the oil and mechanosensory responses to slight changes in air flow as vials switch on and off (Mamiya et al., 2008).

Kenyon cell *in vivo* imaging experiments on empty attp2 control and *mud* RNAi cohorts were done with BZH (Figure 1), as were our previous experiments on HU-ablated animals (Elkahlah et al., 2020). However, due to a technical issue of odor precipitation with BZH, we switched to OCT in imaging experiments in the fall of 2020. The *Tao* RNAi cohort (Figure 3) were done after this date, with OCT. Due to the difficulty of obtaining stable recordings of Kenyon cell populations *in vivo*, in Figure 3 we compare the *Tao* RNAi animals to the empty attp2 controls imaged earlier using BZH (see legend of Figure 3). To ensure that nothing had changed in our imaging setup or the environment between these dates, we imaged a few empty attp2 controls interspersed with *Tao* RNAi animals, using the OCT setup (not shown); we did not observe any striking difference from the control data shown in Figure 3. Indeed, across our functional imaging experiments with Kenyon cells in variously constituted mushroom bodies for the last four years, we have never observed the prevalent and non-selective odor responses that we observe in *Tao* RNAi Kenyon cells.

### Analysis of KC somatic odor responses

We collected data from a total of 14 empty attp2 control animals (14 hemispheres imaged), 13 *mud* RNAi (20 hemispheres imaged), 18 *Tao* RNAi (34 hemispheres imaged), 8 Dscam^3.36.25.1^ animals (17 hemispheres imaged), and 15 yw controls for Dscam (35 hemispheres images). For each animal, the two hemispheres were imaged and analyzed separately. Sometimes, only one hemisphere was imaged due to the preparation and a few hemispheres were excluded from analysis because of poor image quality. For some animals, a second plane of cells was imaged from the same hemisphere if the first one looked unstable or did not have enough cells in view. Mineral oil was delivered first, and then each of four odors were delivered twice, in sequence. Following mineral oil, odor order varied. Given the convention in the field and due to experiments in locusts showing that the first presentation of certain odors can cause distinct Kenyon cell responses to all subsequent presentations of that same odor, we only analyzed the second presentation of each odor (Murthy et al., 2008; Stopfer and Laurent, 1999).

Following functional imaging, we collected a Z-stack of the mushroom body on the two-photon. We used these images to categorize the extent of KC increase in *mud* RNAi animals, by measuring the maximum cross-sectional calyx area. Only sufficiently expanded hemispheres (maximum cross-sectional calyx area > 1800 µm^2^) were included in analysis as “Kenyon cell-increased” animals. While sex differences have not been reported in the mushroom body, we tried our best to include an equal mix of males and females in the imaging. However, after the images were filtered for analysis based on the exclusion criteria provided, we ended up with a disproportionate number of males and females. All samples are accounted for here:

|  | Total hemispheres imaged | Damaged/unresponsive | Poor image quality | Mineral oil frame shaky | Kenyon cells did not look increased | Too much motion | Less than 15 ROIs (cells) stable | Analyzed (# of males and females) |
| --- | --- | --- | --- | --- | --- | --- | --- | --- |
| Empty attp2 control (for RNAi) | 14 | 1 | 2 | 1 | N/A | 4 | 0 | 6 (5 females, 1 male) |
| **mud RNAi** | 20 | 1 | 1 | 2 | 2 | 7 | 1 | 6 (all males) |
| **Tao RNAi** | 34 | 3 | 4 | 0 | N/A | 13 | 2 | 12 (11 females, 1 male) |
| **yw control (for Dscam)** | 35 | 2 | 6 | 0 | N/A | 18 | 1 | 8 (4 females, 4 males) |
| **Dscam<sup>3.36.25.1</sup>** | 17 | 1 | 5 | 0 | N/A | 3 | 1 | 7 (6 females, 1 male) |

All samples were motion-corrected using Suite2p (Pachitariu et al., 2017). Using FIJI “Blind Analysis Tools” package, we blinded ourselves to the motion-corrected datasets. We excluded images if there was too much motion to be able to follow the same cell over time, or the image quality was poor (e.g. warping in the image, cells hard to see). After filtering for these criteria, ROIs were chosen in each of the samples and we measured Δf/f (i.e. (f-f_0_)/f_0_), comparing stimulus frames with pre-stimulus frames. ROIs were manually chosen and we were careful about ensuring that the cell remained within its ROI for all frames. For samples where the ROIs were only stable in 3 out of 4 odors, those images were only used to calculate the proportion of cells that were responsive to each odor, and not in calculating the number of odors each cell responded to. Any image where the mineral oil stimulus frames were not stable was not analyzed, and any image with less than 15 ROIs stable was not analyzed. After these gates, there were 6 empty attp2 controls (for *mud* and *Tao* RNA), 6 *mud* RNAi, 12 *Tao* RNAi, 8 controls (for Dscam), and 7 Dscam^3.36.25.1^ images in the analysis. The baseline fluorescence was determined for each odor delivery by taking the average of the frames within the three seconds before the trigger. The stimulus frame was determined by taking the peak response between 0–4 s after stimulus onset, to account for the valve opening and the two second odor delivery. To define ‘responsive’ cells, we chose to use a cutoff of Δf/f > 0.2 (20% increase in fluorescence over baseline) across samples. This was to avoid obscuring overall differences in responsiveness across the conditions.

For visualization, we used the “Math” and “Image calculator” tools in FIJI to make a custom color heatmap of the Δf/f responses, where cells with values < 0.2 were colored black. The background outside the KC somata was cleared for ease of visualization, and the “Δf/f image” was created by using average value of pixels in baseline frames as f_0_ and average of peak frames as f, and applying the calculation ((f-f_0_)/f_0_) to the images. The maximum value for the color scale varies in each figure due to the diverse range of odor responses in the different circuit manipulations tested. The grayscale image displaying the cells was created by making an average projection of all the frames for that hemisphere.

Control odor responses were in line with what we have previously reported using GCaMP6s and match electrophysiological spiking responses in the previous work by Murthy, but surpass the spiking response rates we observed previously (Elkahlah et al., 2020; Murthy et al., 2008; Turner et al., 2008). Importantly, because GCaMP6s is so sensitive, it is likely to report sub-threshold as well as spiking responses.

### Analysis of MBON odor responses

We collected data from a total of 10 empty attp2 (RNAi control) animals (18 hemispheres imaged), 17 *mud* RNAi animals (29 hemispheres imaged), 11 *Tao* RNAi animals (21 hemispheres imaged), 8 sham animals (15 hemispheres imaged), and 9 HU-treated animals (15 hemispheres imaged). Each hemisphere was analyzed separately. Occasionally, only one hemisphere of the animal was imaged due to the MBON/KC signal not being visible in preparation. Guided by vertical lobe of KCs, we used a single focal plane to image axonal response of γ2,α’1 MBON in each hemisphere. We chose to record axonal responses for the following reasons: (1) axonal imaging is more feasible than dendritic and somatic imaging, as somata of γ2,α’1 MBON are too deep and their GCaMP signal was obscured by fluorescence from the KC lobes, (2) functional KC-MBON connections should allow signal propagation to downstream neurons, which can be seen by axonal activity of MBONs. Similar activity measurements have been used in previous research (Barnstedt et al., 2016). Odor delivery was the same as mentioned in the section above except that all odors were delivered in a certain order.

Following functional imaging, we collected a Z-stack of the mushroom body on the two-photon. We used these images to categorize the extent of KC increase or decrease by measuring the maximum cross-sectional calyx area. All samples are accounted for here:

|  | Total hemispheres | Damaged /unresponsive | KCs did not look increased (mud RNAi) or decreased (ablation) | Too much motion | Too low expression | Analyzed |
| --- | --- | --- | --- | --- | --- | --- |
| **control (attp2)** | 18 | 0 | N/A | 4 | 3 | 11 (9 females, 2 males) |
| **mud RNAi** | 29 | 0 | 6 | 6 | 8 | 9 (1 females, 8 males) |
| **Tao RNAi** | 21 | 0 | N/A | 6 | 3 | 12 (5 females, 7 males) |
| **Sham** | 15 | 1 | N/A | 0 | 0 | 14 (7 females, 7 males) |
| **Ablation** | 15 | 0 | 2 | 2 | 1 | 10 (2 females, 8 males) |

Only sufficiently expanded *mud* RNAi hemispheres (maximum cross-sectional calyx area > 2200 µm^2^; for control(attp2) and *mud* RNAi, each calyx was imaged by two-photon right after Calcium imaging) were included in the analysis of the KC-increased animals. Only sufficiently HU-ablated hemispheres (maximum cross-sectional calyx area < 2100 µm^2^; for Sham and Ablation, each calyx was imaged in post hoc dissected brains by two-photon). For RNAi modulation, as LexAop-GCaMP6s in all animals are heterozygous, we excluded images where axonal baseline fluorescence was not distinguishable from the background; these are categorized as “Too low expression”. In sham and KC-reduced condition, LexAop-GCaMP6s is homozygous and provided robust axonal GCaMP expression. All samples were motion-corrected in batch using Suite2p (Pachitariu et al., 2017). ROIs were automatically chosen by applying triangle auto-threshold method in FIJI (v2.3.0) to find pixels with higher intensity than the background in the axonal area, including axon terminals and fibers. For Δf/f, we chose the time window 5 s to 2 s before onset of odor stimulus as baseline, while peak responses were the maximum response between 0 s to 12 s after odor stimulus onset. Corresponding visualization was performed similar to the method described above by averaging 5s to 2s before onset of odor as baseline, and averaging 2s after maximum response as peak. There is no “0.2 cut-off”.

### **Y-** arena

Single fly behavior experiments were performed in a novel olfactory Y-arena. A detailed description of the apparatus is provided in (Rajagopalan et al., 2022). Briefly, the Y chamber consists of two layers of white opaque plastic. The bottom is a single continuous circular layer and serves as the floor of the Y that flies navigate. The top is a circular layer with a Y shaped hole serving as the walls of the Y. The length of each arm from center to tip is 5 cm and the width of each arm is 1 cm. These two layers are placed underneath an annulus of black aluminum. A transparent glass disk is located in the center of this annulus and acts as the ceiling of the Y -allowing for video recording of experiments. This transparent disk is rotatable and contains a small hole used to load flies. The black annulus houses three clamps that hold the circular disk in place. All three layers are held together and made airtight with the help of 12 screws that connect the layers. The Y chamber is mounted above an LED board that provides infrared illumination to monitor the fly’s movements, and red (617 nm) light for optogenetic activation. Experiments were recorded from above the Y using a single USB3 camera (Flea3, model: FL3-U3-13E4M-C: 1.3 MP, 60 FPS, e2v EV76C560, Mono; Teledyne FLIR, with longpass filter of 800 nm). Each arm of the Y has a corresponding odor delivery system, capable of delivering up to 5 odors. For our experiments, olfactometers injected air/odor streams into each arm at a flow rate of 100 ml/min. The center of the Y contains an exhaust port connected to a vacuum, which was set at 300 ml/min using a flow meter (Dwyer, Series VF Visi-Float® acrylic flowmeter) -matching total input flow in our experiments. We wrote custom MATLAB code (MATLAB 2018b, Mathworks) to control the Y-arena and run experiments.

### Behavioral Protocol

*Odorant information:*

The following odorants were used to form cue-reward relationships:

1. 3-Octanol (OCT). *Y-arena:* diluted in paraffin oil at a 1:500 concentration and then air-diluted to a fourth of this concentration.
2. 4-Methyl-cyclo-hexanol (MCH). *Y-arena:* diluted in paraffin oil at a 1:500 concentration and then air-diluted to a fourth of this concentration.

*Y-arena behavioral task structure and design:*

Flies were inserted randomly into one of the three arms. This arm was injected with a clean airstream and the two odors were randomly assigned to the other two arms. For a given fly, one of the odors was paired with reward 100% of the time. Once a fly reached the choice zone of either odor arm a choice was considered to have been made. If the rewarded odor was chosen, the fly was rewarded with a 500ms flash of red LED (617 nm, 1.9mW/cm^2^) to activate the appropriate reward-related neurons. The arena was then reset with the arm chosen by the fly injected with clean air and odors randomly assigned to the other two arms. This was repeated for many trials with flies making either 40 or 60 naive choices where neither option was rewarding and 40 or 60 training choices where one option was consistently rewarded.

For each behavior batch, at least one control fly was also run the same day to account for any batch effects. The exception to this is the batch of *Tao* RNAi flies. Due to technical reasons, we were unable to get behavior results from control flies on the same day. Therefore, we used the controls from the *mud* RNAi behavior batches to compare their behavior to, since the control genotype is the same for both the RNAi conditions.

## QUANTIFICATION AND STATISTICAL ANALYSIS

### Image analysis

We analyzed males for immunohistochemistry in our manipulations. For functional imaging experiments, we used mixed-sex populations, and did not observe any correlation with sex (not shown). The Y-arena behavior used females due to the size of the arena not being optimal for males. Therefore, the flies dissected post-behavior and used for calyx area quantification in those animals and Brp staining (Figures 1D, 2H, 6F) were also females. Sex differences in the fly are well-documented, including in our own previous work, and anatomic and physiologic sex differences have not been observed in the mushroom body (Brovkina et al., 2021; Clowney et al., 2015). Any brains that appeared damaged from dissections, or those with the mushroom body region obscured due to insufficient tracheal removal, were not included in the analysis.

Researchers performing quantification could not generally be blinded to experimental condition due to the overt changes in neuron numbers and brain structures induced by our manipulations. However, analysis was performed blind to the goals of the experiment when possible, and quantitation of features on the anterior and posterior sides of the brain were recorded independent of one another and merged after all quantifications were completed.

Moreover, many of our analyses make use of variation within an experimental condition or genotype, providing an additional bulwark against observational bias.

### Calyx area

To measure the size of the mushroom body calyx, we used genetically-encoded fluorescence driven in Kenyon cells by OK107-Gal4. In cases where the fluorescent marker was not added, we used markers such as ChAT to visualize the structure. We then identified its largest extent in Z (i.e. along the A-P axis), outlined it in FIJI (as in white outlines in Figure 2H) and calculated the cross-sectional area using the ‘Measure’ command. We have previously shown the calyx area to positively correlate with KC number and hence, serve as a readout for KC number in the hydroxyurea-treated, KC-reduced and *mud* RNAi-driven, KC-increased conditions (Elkahlah et al., 2020).

### Kenyon cell numbers

To count Kenyon cells, we again used genetically-encoded fluorescence driven by OK107-Gal4. We counted labeled somata in every third slice in the stack (every third micron along the A-P axis), with reference to DAPI to distinguish individual cells from one another. We initially determined that somata in slice 0 could also be seen in slices −2, –1, +1, and +2 but not in slice −3 or +3. To avoid double-counting, we therefore counted every third micron. For Figure 4, Kenyon cells were counted using a cellular segmentation tool called Cellpose (Stringer et al., 2021). The confocal stack for each hemisphere was split into single planes every third micron, and those slices were cropped to where the KC somata are present and given as input image into Cellpose. The count from each slice was then summed up to get the total count. The software used the Kenyon cell fluorescence, nuclear signal (DAPI) in the Kenyon cells, and an automatically calibrated cell diameter to identify individual cells. We verified that the counts obtained matched manual counts.

### Kenyon cell and projection neuron neuroblast state

For quantifying Kenyon cell neuroblast state in Figure 6, as we have done previously (Elkahlah et al., 2020), we used 58F02 to fluorescently label late-born Kenyon cells and counted clumps of labeled somata surrounding the calyx as well as groups of labeled neurites leaving the calyx and entering the pedunculus. These estimates usually matched; in the few cases where they did not, we used the number of axon clumps, as somata are closer to the surface of the brain and more susceptible to mechanical disruption during dissection. This resulted in numbers between 0-4 for each hemisphere. Any additional labeling of neurons by 58F02 was easy to discriminate from KCs as those neurons did not enter the calyx or pedunculus.

For scoring presence of the PN neuroblast (PN lNB/BAlc) in sham-treated and hydroxyurea-treated animals, we included Mz19 driving a fluorescent reporter in PNs from this lineage (Ito et al., 1998; Jefferis et al., 2004). In the absence of lNB/BAlc, 12 glomeruli lose their typical PN partners; 40 glomeruli are innervated by the anterodorsal PN neuroblast, which is not affected by HU ablation (Jefferis et al., 2001; Yu et al., 2010). Mz19 labels PNs that innervate DA1 and VA1d glomeruli on the anterior side, and DC3 on the posterior side of the antennal lobe; DA1 is innervated by lNB/BAlc PNs (Jefferis et al., 2004). We quantified the presence or absence of the most distinctly labeled glomerulus – DA1, as a way to score the PN lNB/BAlc getting ablated. In some cases, we observed mislocalization of DA1. For simplification, we scored this in the “absence” category.

### Projection neuron bouton numbers

To count aggregate boutons, we used ChAT signal as previously described in (Elkahlah et al., 2020). We counted as separate structures ChAT signals that were compact and appeared distinct from one another and that were 2+ micrometers in diameter (except in Figure 2, *Tao* RNAi images where boutons appeared smaller than wildtype). When KC fluorescence was available, two boutons would be counted separately if ChAT signals were separated by KC signal. As with KC somata, we found that boutons in slice 0 often appeared in slices −1 and +1 as well, but never in slices −2 or +2. To avoid overcounting, we began at the most superficial slice in the stack where boutons were visible, and counted every other slice, i.e. every second micron. For ChAT^+^ counts in Figure 4, we combined data from two batches of experiments done on different dates – one of them carried a KC-driven fluorescent marker, the other batch did not.

### Kenyon cell claw counts

To count Kenyon cell claws, we used sparse labeling of Kenyon cells driven by mBitbow2.2. We traced the dendritic projections by scrolling through the z stack, counting any claw-like dendritic structures that appeared and stopping when we reached the pedunculus. To ensure we counted all the cells labeled, we matched the puncta-like signal in the pedunculus to the number of KC soma visible. When ChAT or nc82 signal was available, we used that to define the calyx extent. We generally found the mAmetrine Bitbow signal (shown in Figure 2C) to be the cleanest and brightest for counting the claws. We then counted the number of KC soma labeled in the same color in which the claws were counted to obtain a “claws per KC” measurement by dividing claws of that color by the KC soma labeled in that color. Any image with more than 6 cells labeled was not counted as it was considered too dense to get an accurate claw count. An exception to this were the Dscam^3.36.25.1^ animals (Figure 4) where the drastic claw number decrease allowed us to count claws for up to 10 Kenyon cells.

### Bruchpilot density

We measured Bruchpilot intensity in the calyx as a readout of synaptic density. First, we identified the Z plane with the largest extent of the calyx in the A-P axis, and then took three measurements of the average fluorescence signal in the Brp channel in a defined ROI region measuring ∼40 μm^2^. The three measurements were taken randomly in different locations in the calyx to account for any variability in intensity. For normalization, the calyx Brp signal was divided by Brp signal measured in an unmanipulated brain region, the protocerebral bridge.

We chose this brain region as it was the closest to the calyx and was in the field of view in all calyx images taken. These quantifications were done in the “63X hi-res-1“ and “63X hi-res-2” sets of images (defined in the “Immunostainings” section above).

### Correlation matrix of odor responses

To assess the linear relation between each pairwise odor comparison across the cells in the control and *Tao* RNAi animals, we plotted a Pearson correlation matrix using the “corrplot” package in R (v3.5.1). The range of the Pearson correlation is –1 to +1, with positive values indicating a positive correlation and negative values indicating a negative correlation.

### Behavior analysis

Analysis of data resulting from the Y-arena was performed using MATLAB 2020b (Mathworks). In a given experiment, the *(x,y)* coordinates of the fly and metadata from the experiment to determine arm-odor relationships were analyzed to determine which odor the fly chose on a given trial. This allowed us to make a list of choices made by the fly as a function of trial number. This list was used to calculate the fly’s average preferences for the rewarded odor over the unrewarded odor (in the naïve block, this was defined based on which odor was rewarding in the training block). Time spent in the air arms was not recorded as a choice.

The proportion correct choice metric was calculated for both naïve and training blocks and used in Figures 6 and 7. This was defined as the number of choices to the rewarded option divided by the total number of choices. Only the second half of trials in each block were used. The first half of trials were excluded from this calculation as flies take around 15-20 trials to reach a plateau learnt performance in the training block and we did not want this metric to be diluted by trials in which learning was incomplete.

To plot individual learning curves plotted in Figures 6 and 7, the cumulative number of choices towards the rewarded and unrewarded options were calculated as a function of trial number and then plotted against each other.

Since the control genotype for *mud* RNAi, KC-increased and *Tao* RNAi, KC claw-increased animals was the same, we plotted the behavioral data from these three genotypes together.

### Statistical considerations

Determination of sample size: Brains were prepared for imaging in batches of 5–10. In initial batches, we assessed the variability of the manipulation, for example if we were trying to change Kenyon cell number, we looked at how variable the size of the Kenyon cell population was following the manipulation. We used this variability to determine how many batches to analyze so as to obtain enough informative samples. To avoid introducing statistical bias, we did not analyze the functional consequence of the manipulation until after completing all batches; for example, if the manipulation was intended to alter Kenyon cell claw numbers, we did not quantify odor responses until after completing all samples. Similarly, we did not measure the effect on other cell types (such as assessing projection neuron bouton phenotypes) until after completion of all samples. Genotypes or conditions being compared with one another were always prepared for staining together and imaged interspersed with one another to equalize batch effects except in occasional cases that have been highlighted in methods above.

Since our developmental manipulations seemed to affect the mushroom body in each hemisphere independently and variably, we treated each hemisphere as an independent sample in our staining and functional imaging. In the case of hydroxyurea-treated animals, if one hemisphere was affected severely, the other one is likely affected to a similar extent but not necessarily equally, e.g. we observed KC clone counts of “[1,0]” or “[3,2]” but never “[4,0]”.

Criteria for exclusion, treatment of outliers: We excluded from analysis samples with overt physical damage to the cells or structures being measured. In figures and analyses, we treated outliers the same way as other data points. For *in vivo* functional imaging experiments, full criteria for inclusion and exclusion of each sample are discussed above under “Analysis of KC somatic odor responses” and “Analysis of MBON odor responses”.

Statistical tests applied are mentioned in each figure legend along with the p-value significance. To communicate our findings in the simplest and most complete way, we have displayed each data point for each sample to allow readers to assess effect size and significance directly. When sample size could be determined from the figures, we did not explicitly state it in the figure legends.

All statistical analysis were performed in GraphPad Prism, Excel, or R.

## Author Contributions

MA and EJC led the project design and coordinated all authors. MA performed genetic discovery experiments, anatomic analyses, and Kenyon cell functional imaging with the assistance of DLW. AER designed and performed behavioral analyses with MA under the supervision of GCT. YP designed and performed MBON experiments. YL and DC designed and created the mBitbow 2.2 system. MA led MT and EAP in analysis of functional and anatomic data. The Janelia Project Technical Resources team led by Gudrun Ihrke performed brain dissections of flies, histological preparations (KCC and Yisheng He), and confocal imaging (CPC) for behavioral analyses. AP improved and implemented HU ablation methods. MA, EJC, and YP wrote the original manuscript with contributions from YL and AER. All authors revised the manuscript. Authors MA and EAP disagree that coffee is superior to tea, AER and DLW do not touch either, and while YP enjoys coffee and tea just fine, he actually prefers Diet Coke.

## Acknowledgments

We thank the Bloomington Drosophila Stock Center, and Bing Ye and Tzumin Lee labs for providing fly strains; the MCDB Imaging Facility and Gregg Sobocinski for technical assistance with confocal microscopy; the Janelia Project Technical Resources team led by Gudrun Ihrke for brain dissections, histology and confocal imaging for behavior experiments; and members of the Clowney lab for comments on the manuscript. Funding was provided by NIDCD R01DC018032 and Sloan Research Fellowship to EJC; Janelia Visiting Scientist Program and Rackham Predoctoral Fellowship to MA; NIMH R01MH110932 to DC; and HHMI to GCT. EJC is the Rita Allen Foundation Milton Cassel Scholar, a McKnight Scholar, and a Pew Biomedical Scholar.

---

**Figure S1.**
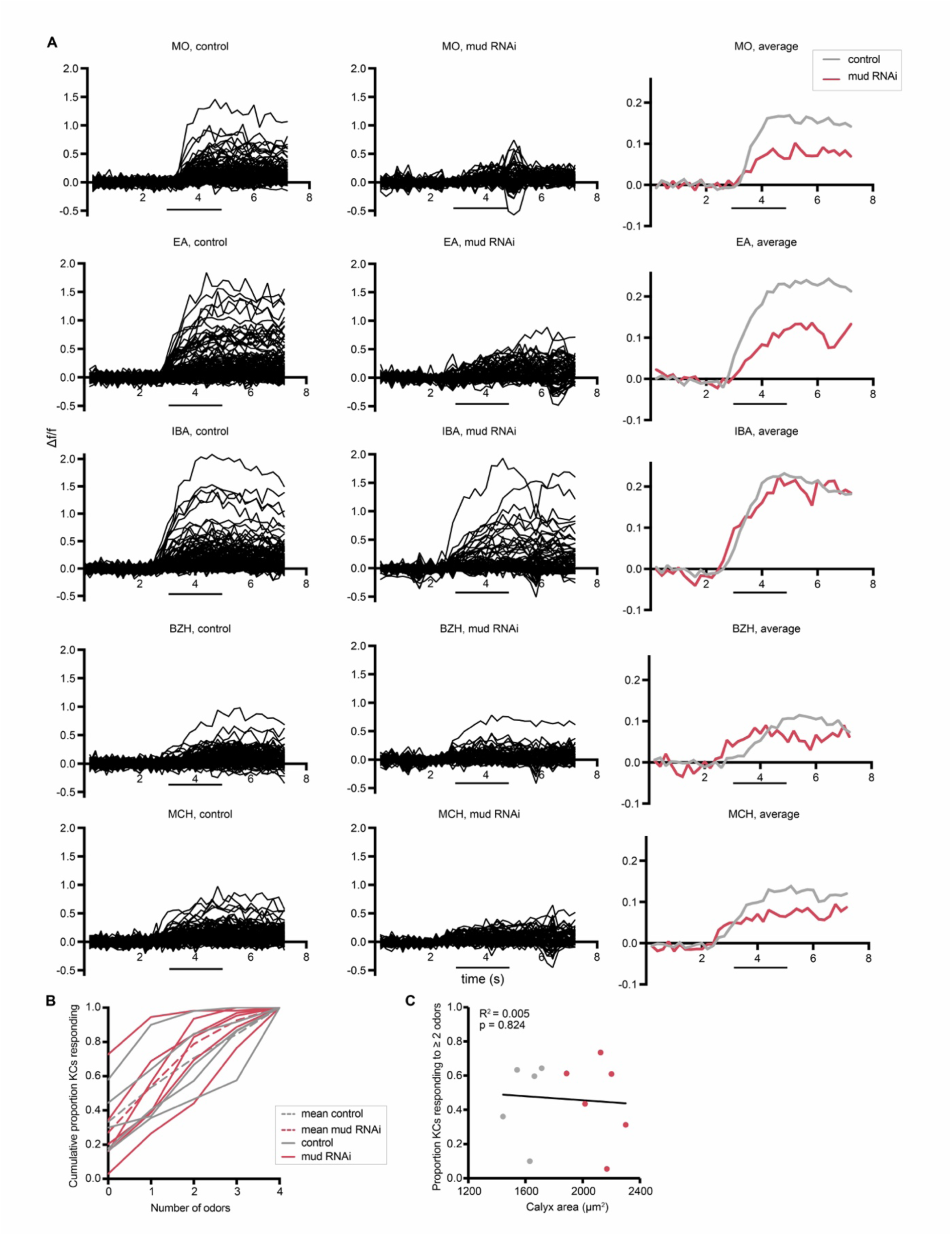
Supplemental data related to Figure 1. (A) Odor responses over time for Kenyon cells shown in Figure 1G. x axes show seconds, y axes show Δf/f. Black bars indicate odor delivery. Each black line is one cell, with graphs at right showing responses averaged across all cells of the sample. Each cell was normalized to average fluorescence in the 3 s period before stimulus onset. MO: Mineral oil (mechanosensory and solvent control), EA: Ethyl Acetate, IBA: Isobutyl Acetate, BZH: Benzaldehyde, MCH: Methylcyclohexanol. An example of a motion artifact can be seen in ‘MO, mud RNAi’ trace around 5s. n = 124 cells (control), 62 cells (*mud* RNAi). (B) Cumulative proportion of Kenyon cells responding from 0 up to 4 odors. Each line represents an individual control hemisphere (gray) or increased-KC *mud* RNAi hemisphere (red), with the mean of all control or *mud* RNAi samples shown with a dotted line. (C) Relation of proportion of KCs responding to 2 or more odors, and maximum cross-sectional calyx area of control (gray) and KC-increased *mud* RNAi (red) hemispheres. Here and throughout, linear regressions are performed across all data points shown in the figure, i.e., for the distribution of the two sample types taken together across the variation in calyx size.

**Figure S2.**
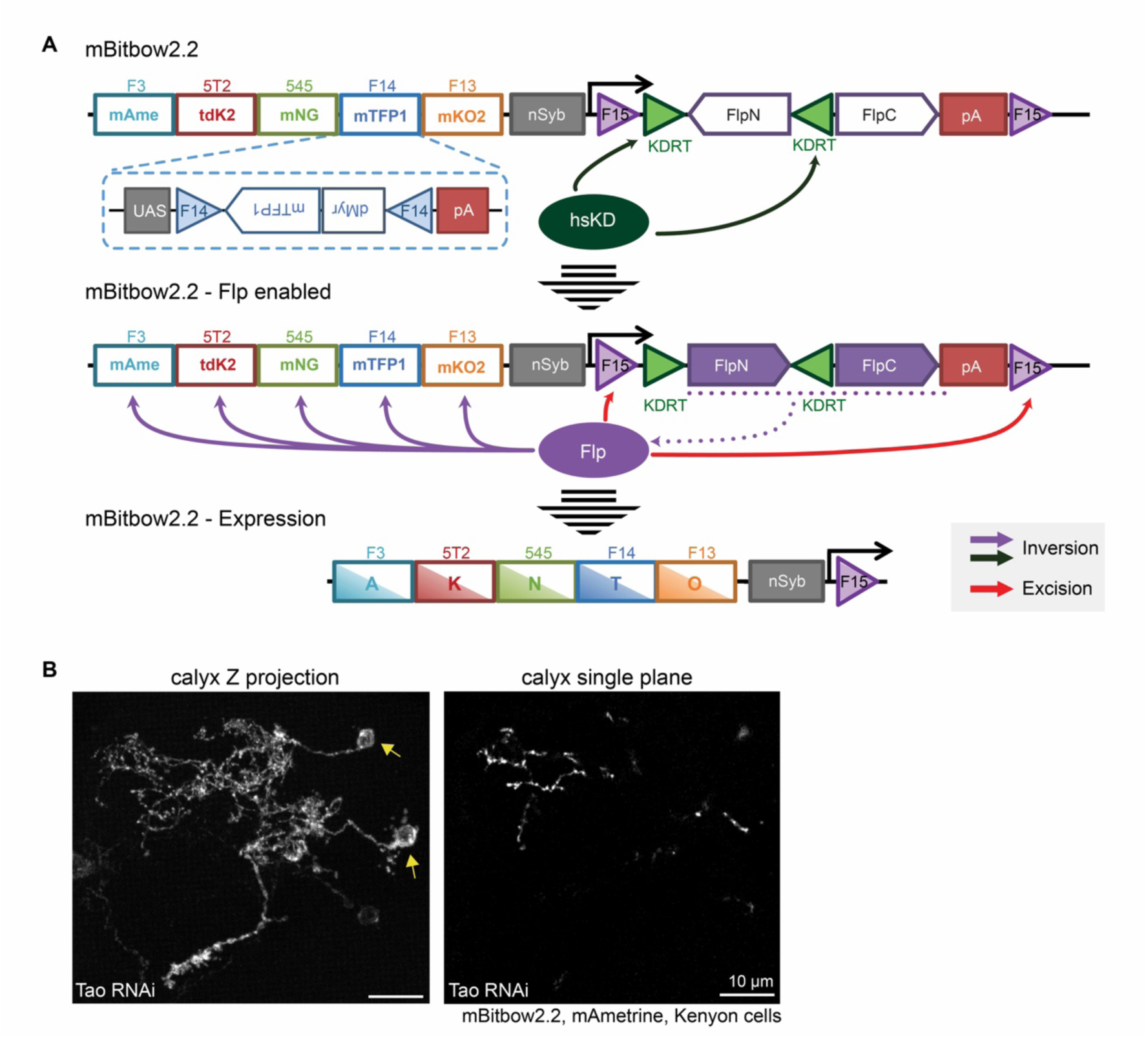
mBitbow2.2 schematic and dendritic morphology in *Tao*-knockdown Kenyon cells. (A) Detailed schematic of mBitbow2.2 design; a simplified version is shown in Figure 2A. (B) Images of sparsely labeled KCs in a KC>*Tao* RNAi calyx showing highly branched but non-clawed dendritic projections that we observe in some samples: maximum z projection (left), single confocal slice (right). In the example shown, arrows mark 2 KCs labeled by mBitbow2.2 mAmetrine. Faint third soma is labeled by an additional mBitbow2.2 color.

**Figure S3.**
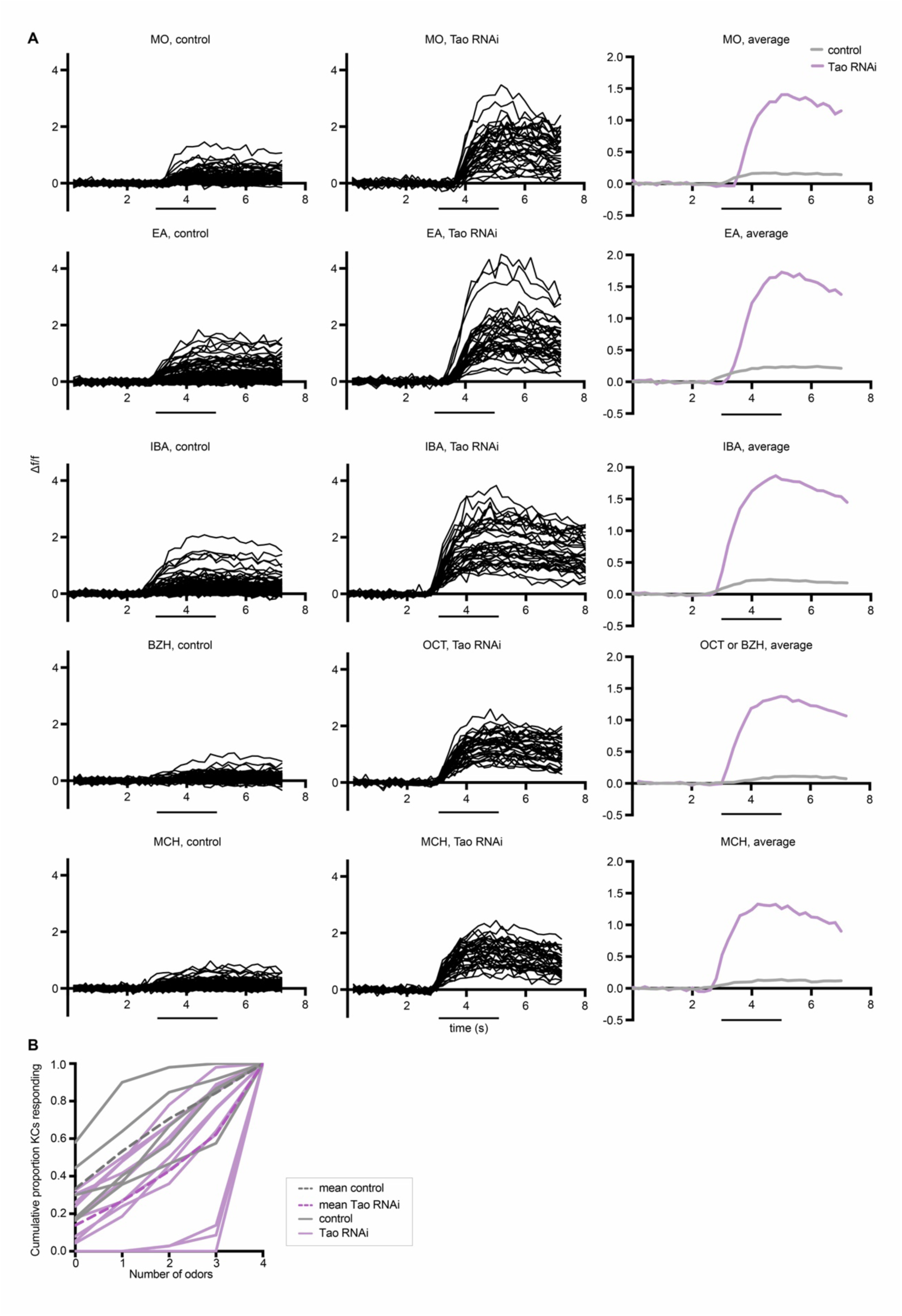
Odor responses over time for Kenyon cells shown in Figure 3A. (A) x axes show seconds, y axes show Δf/f. Black bars indicate odor delivery. Each black line is one cell, with graphs at right showing responses averaged across all cells of the sample. Each cell was normalized to average fluorescence in the 3 s period before stimulus onset. MO: Mineral oil (mechanosensory control), EA: Ethyl Acetate, IBA: Isobutyl Acetate, BZH: benzaldehyde, OCT: Octanol, MCH: Methylcyclohexanol. Control sample is the same sample shown in Figure S1; we have replotted the data to allow quantitative comparison with the robust responses of excess-claw KCs. n = 124 cells (control), 36 cells (*Tao* RNAi). (B) Cumulative proportion of Kenyon cells responding from 0 up to 4 odors. Each line represents an individual control hemisphere (gray) or *Tao* RNAi hemisphere (purple), with the mean of all control or *Tao* RNAi samples shown with a dotted line.

**Figure S5.**
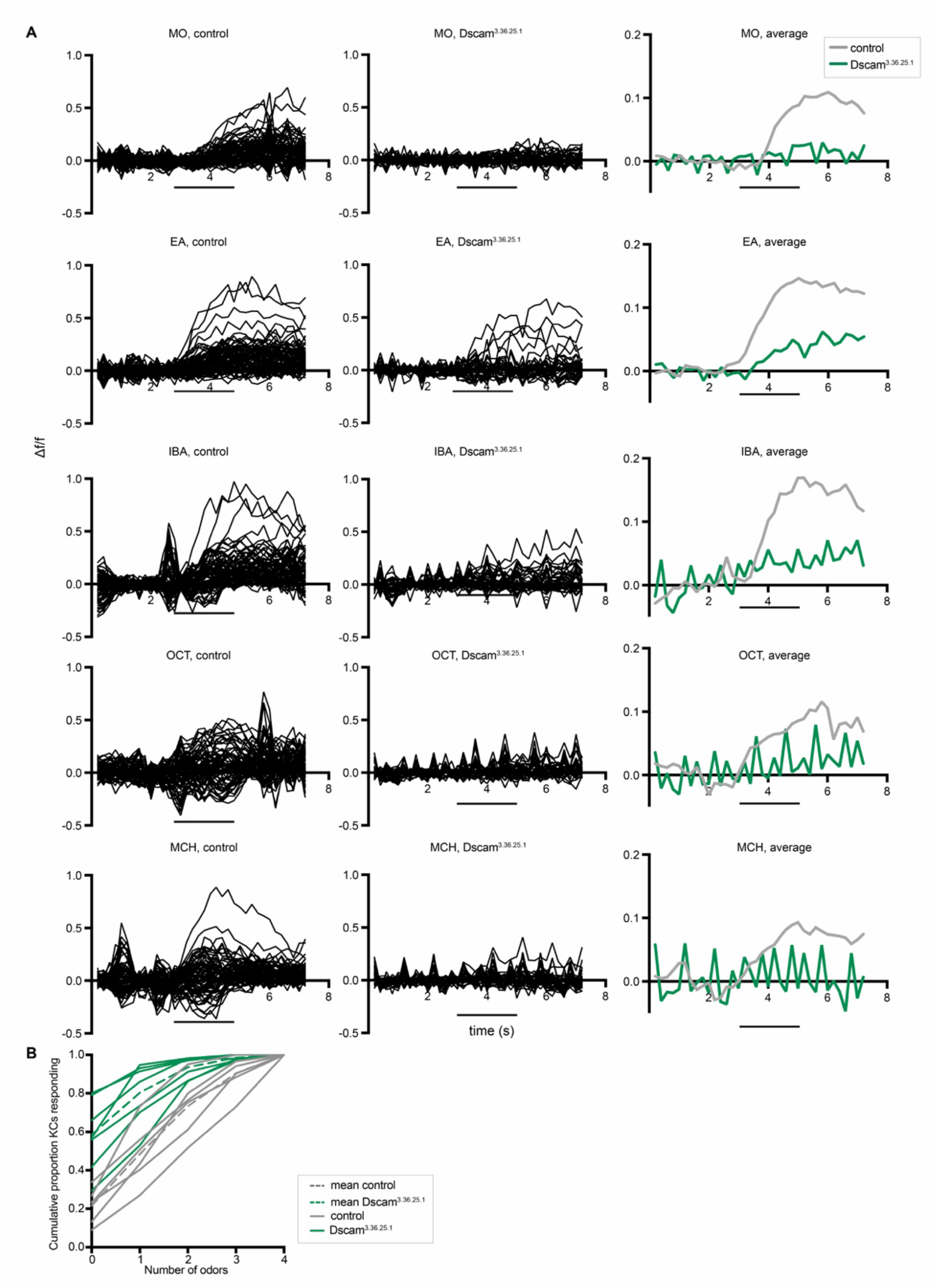
Odor responses over time for Kenyon cells shown in Figure 5A. (A) x axes show seconds, y axes show Δf/f. Black bars indicate odor delivery. Each black line is one cell, with graphs at right showing responses averaged across all cells of the sample. Each cell was normalized to average fluorescence in the 3 s period before stimulus onset. MO: Mineral oil (mechanosensory control), EA: Ethyl Acetate, IBA: Isobutyl Acetate, OCT: Octanol, MCH: Methylcyclohexanol. n = 72 cells (control), 35 cells (Dscam^3.36.25.1^). (B) Cumulative proportion of Kenyon cells responding from 0 up to 4 odors. Each line represents an individual control hemisphere (gray) or Dscam^3.36.25.1^ hemisphere (green), with the mean of all control or Dscam^3.36.25.1^ samples shown with a dotted line.

**Figure S6.**
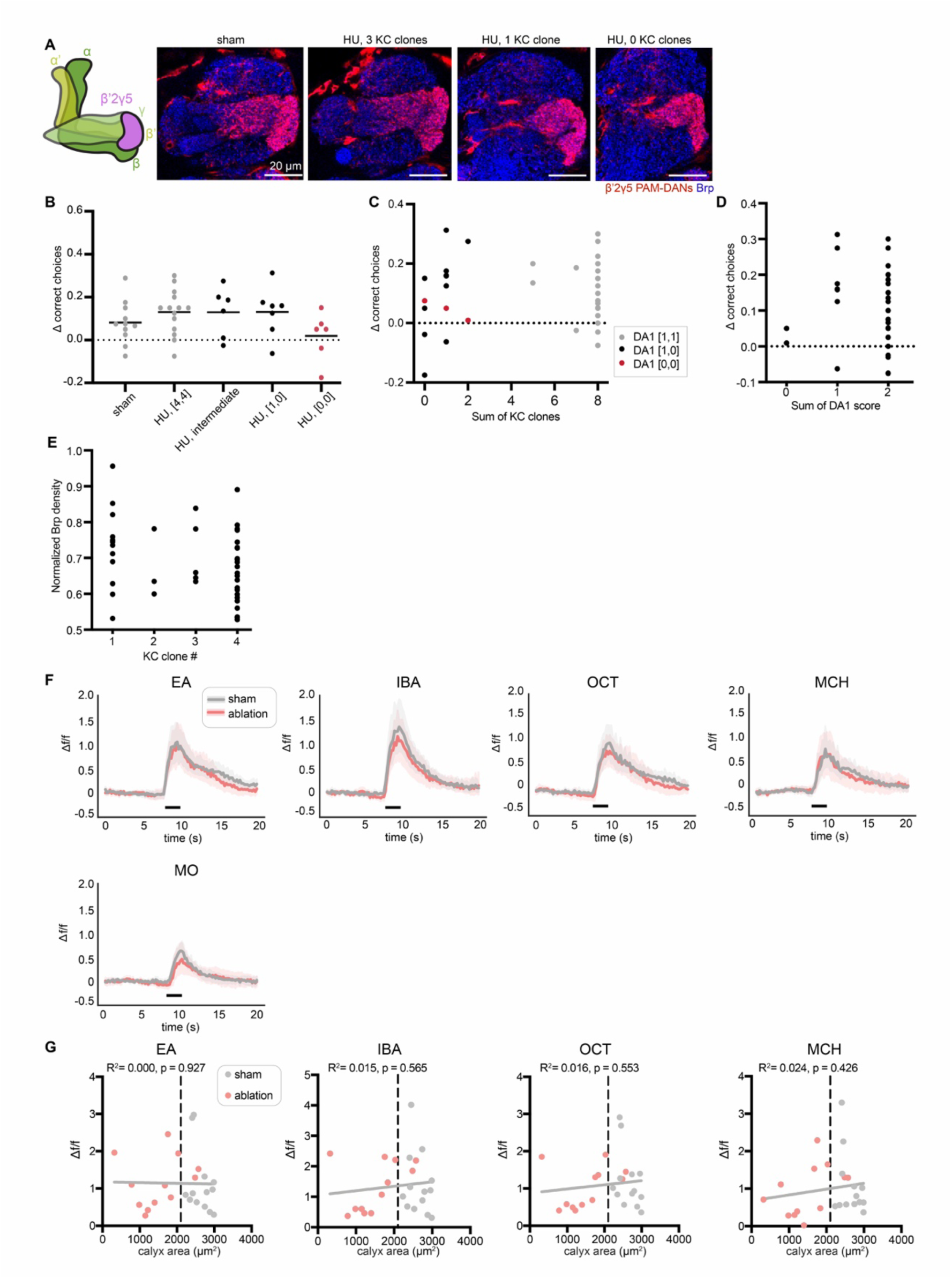
Supplemental data related to Figure 6. (A) Left: Schematic of the mushroom body lobe anatomy with KCs in green and β’2γ5 PAM-DANs in magenta. Axons of β’2γ5 DANs in the lobe compartment are shown. Right: Single confocal slices of the MB lobe (identified by location and Brp staining shown in blue). Mz19 driver labels β’2γ5 DANs (red). Representative images shown of sham-treated and HU-treated hemispheres with 3, 1, or 0 KC clones. (B) Δ correct choices of sham-treated and HU-treated animals shown in Figure 6D. Black bars indicate the medians. In B-D, each data point is an individual fly. (C) Relation of Δ correct choices, sum of KC clone number from both hemispheres and lNB/BAlc ablation status. DA1 present in both hemispheres is indicated as “DA1 [1,1]”, presence in one hemisphere is indicated as “DA1 [1,0]”, and absent in both hemispheres is indicated as “DA1 [0,0]” . (D) Relation of Δ correct choices to sum of DA1 score (presence/absence). The data shown excludes fully KC ablated animals. Jitter added in (C-E) to display all the data points. (E) Relation of normalized Brp density in the mushroom body calyx to Kenyon cell clone number in sham-treated and HU-treated animals, excluding fully ablated animals as there is no calyx present. (F) Average odor responses over time for γ2,α’1 MBONs in Figure 6I. x axes show seconds, y axes show Δf/f. Black bars indicate odor delivery. Shadows are 95% confidence intervals for corresponding averaged traces. Each cell was normalized to average fluorescence in the 5 s to 2 s period before stimulus onset. MO: Mineral oil (mechanosensory control), EA: Ethyl Acetate, IBA: Isobutyl Acetate, OCT: Octanol, MCH: Methylcyclohexanol. n= 12 hemispheres (sham), 10 hemispheres (ablation). Only HU-partially ablated hemispheres smaller than every control (maximum cross-sectional calyx area < 2100 µm^2^) are included. This cutoff is labeled as black vertical dashed line in (G). (G) Relationship between γ2,α’1 MBON peak odor responses and maximum cross-sectional calyx area. Gray line is linear regression for all samples. n= 12 hemispheres (sham), 12 hemispheres (ablation). 2 HU-treated animals with maximum cross-sectional calyx area > 2100 µm^2^ are also shown.

**Figure S7.**
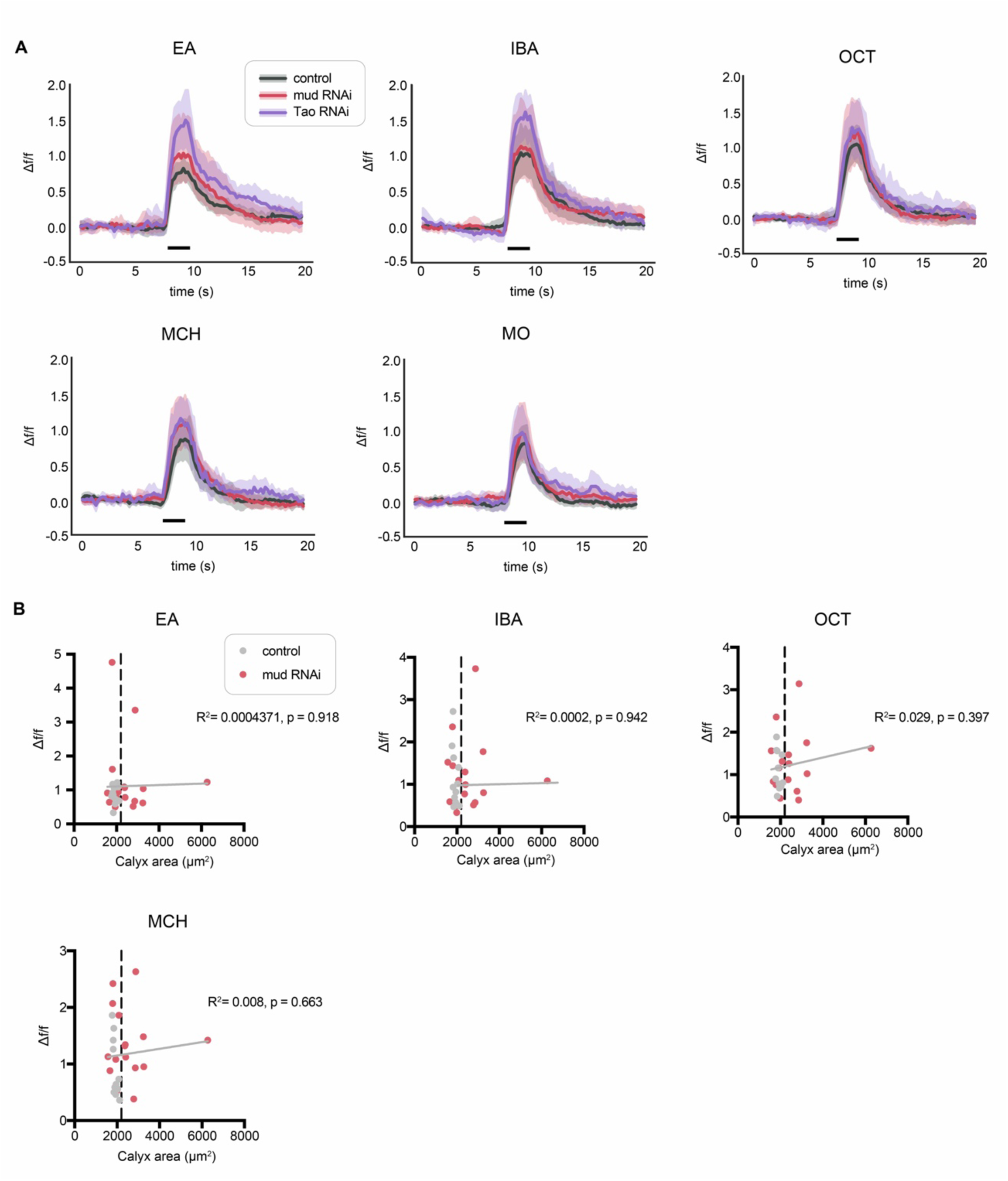
Odor responses for γ2,α’1 MBON in *mud* RNAi and *Tao* RNAi animals. (A) Average odor responses over time for γ2,α’1 MBONs shown in Figure 7G. x axes show seconds, y axes show Δf/f. Black bars indicate odor delivery. Shadows are 95% confidence intervals for corresponding average trace. Each cell was normalized to average fluorescence in the 5 s to 2 s period before stimulus onset. MO: Mineral oil (mechanosensory control), EA: Ethyl Acetate, IBA: Isobutyl Acetate, OCT: Octanol, MCH: Methylcyclohexanol. n= 10 hemispheres (control), 9 hemispheres (*mud* RNAi), 12 (*Tao* RNAi). For *mud* RNAi, only Kenyon cell-increased hemispheres (maximum cross-sectional calyx area > 2200 µm^2^) are included. This threshold is labeled as black vertical dashed line in (B). (B) Relationship between γ2,α’1 MBON peak odor responses and maximum cross-sectional calyx area. Gray line is linear regression for all samples. n= 10 hemispheres (control), 15 hemispheres (*mud* RNAi); six hemispheres from *mud* RNAi calyces with calyx cross-sectional area overlapping controls (< 2200 µm^2^) are included among these 15.

**Supplemental Table 1:** Quantitative variables of mushroom body calyx wiring and function

| Parameter | MB Name | Variable Name | Number in natural MB | Number engineered here |
| --- | --- | --- | --- | --- |
| Number of inputs | Olfactory projection neuron types | N | 52 | 40-52 |
| Number of expansion layer neurons | Kenyon cells per hemisphere | M | 2000 | 500-4000 |
| Expansion ratio (M/N) | Kenyon cell number/odor channels | E | 38 | 10-77 |
| Number of inputs to each expansion layer neuron | Claw number | K | 5 | 1-12 |
| Spiking threshold | Number of input channels active for KC to spike | ? | Usually 2 or more, occasionally 1 | *We suspect it has not changed |
| Strength of feedback inhibition | APL activity | Sigma | Unknown | Unknown |
| Total connections between sensory and expansion layer | Total number of Kenyon cell claws | S | 10,000 | 2,000-24,000 |

**Supplemental Table 2:** Summary of the effects of our developmental manipulations on mushroom body calyx connectivity variables

| Condition | Result | N | M | E | K | KC spike threshold | Sigma | S |
| --- | --- | --- | --- | --- | --- | --- | --- | --- |
| Wild type | KCs respond to 0-1 odors | 52 | 2000 | 38 | 5-6 | ~2 claws | unknown | 10,000 |
| 5HU ablation | Fewer KCs, but each responds to the same number of odors | 40-52 | 500-2000 | 10-50 | 5-6 | ~2 claws (inferred) | Proportional to KCs active (inferred) | 2,500-10,000 |
| Mud knockdown | More KCs, but each responds to the same number of odors | 52 | 2000-4000 | 38-77 | 6 | ~2 claws (inferred) | Proportional to KCs active (inferred) | 10,000-24,000 |
| Tao knockdown | Each KC responds to more odors | 52 | 1700 | 32 | 12 | ~2 claws (inferred) | Proportional to KCs active (inferred) | ~20,400 |
| Dscam[TM1] overexpression | Each KC responds to fewer odors | 52 | 1800 | 35 | 1 | 1-2 claws (inferred) | Proportional to KCs active (inferred) | 1800 |

